# Complementary MR measures of white matter and their relation to cardiovascular health and cognition

**DOI:** 10.1101/2025.02.08.637219

**Authors:** Petar P. Raykov, Marta Correia, Kamen Tsvetanov, Rafael N. Henriques, Alberto Del Cerro León, Matthew Bracher-Smith, Valentina Escott-Price, Yordan P. Raykov, Richard N. Henson

## Abstract

Magnetic Resonance Imaging (MRI) offers many ways to non-invasively estimate the properties of white matter (WM) in the brain. In addition to the various metrics derived from diffusion-weighted MRI, one can estimate total WM volume from T1-weighted MRI, WM hyper-intensities from T2-weighted MRI, myelination from the T1:T2 ratio, or from the magnetisation-transfer ratio (MTR). Here we utilise the presence of all of these MR contrasts in a population based life-span cohort of 650 healthy adults [[CamCAN cohort](https://cam-can.mrc-cbu.cam.ac.uk/)] to identify the latent factors underlying the covariance of 11 commonly-used WM metrics. Four factors were needed to explain 89% of the variance, which we interpreted in terms of 1) fibre density / myelination, 2) free-water / tissue damage, 3) fibre-crossing complexity and 4) microstructural complexity. These factors showed distinct effects of age and sex. To test the validity of these factors, we related them to measures of cardiovascular health and cognitive performance. Specifically, we ran path analyses 1) linking cardio-vascular measures to the WM factors, given the idea that WM health is related to cardiovascular health, and 2) linking the WM factors to cognitive measure, given the idea that WM health is important for cognition. Even after adjusting for age, we found that a vascular factor related to pulse pressure predicted the WM factor capturing free-water / tissue damage, and that several WM factors made unique predictions for fluid intelligence and processing speed. Our results show that there is both complementary and redundant information across common MR measures of WM, and their underlying latent factors may be useful for pinpointing the differential causes and contributions of white matter health in healthy aging.

## Introduction

The integrity of white matter (WM) in the human brain is known to decline with age, and this decline is thought to contribute to the age-related declines in cognitive performance, particularly fluid abilities like processing speed and episodic memory (Gunning-Dixon et al., 2009; Hernández et al., 2013; Lawrence et al., 2021; Mendez Colmenares et al., 2023; Moon et al., 2018; Prins & Scheltens, 2015; Schilling et al., 2023). However, WM properties can be measured in many ways, such as the magnetic resonance imaging (MRI) contrasts of T1-weighting (T1w), T2-weighting (T2w), magnetisation transfer-weighting (MT) and diffusion-weighting (DWI). Researchers may often focus on a single WM measure, partly to avoid a multiple comparison problem, but this means that often is difficult to understand the unique and synergistic contribution to health and cognition of different WM measures. Here we utilised the range of such MRI contrasts available within a cohort of healthy adults aged 18-89 to answer two main questions: 1) How many factors can capture the variance in white matter measures, and 2) How do these factors relate to cardiovascular health and cognition?

Ageing is well-known to reduce the total volume of WM in the last decades of life (in addition to reducing gray-matter volume; Pannese, 2011). These white matter volume (WMV) reductions could reflect the degree of axonal degeneration or demyelination (Wozniak & Lim, 2006), and can be estimated by segmenting T1w images using automated techniques. Early MRI studies also observed bright regions on T2w images, which tend to become larger and more numerous with age. The prevalence of these “white matter hyper-intensities” (WMHI) has been associated with impaired cognition and increased risk of neurodegenerative disease (Botz et al., 2023; d’Arbeloff et al., 2019, 2021; Debette & Markus, 2010; Fuhrmann et al., 2019; Torres-Simon et al., 2023, 2024). They are thought to reflect macroscopic damage to the WM, often due to vascular pathology.

Other MR sequences have been developed to provide more direct measures of myelin content. For instance, magnetisation-transfer-weighting (MTw) relies on the magnetisation exchange between macromolecule-bound protons (in myelin, for example) and free water protons. The amount of magnetisation transferred between the two pools of protons can be quantified by computing the ratio between two images: one with and one without a RF saturation pulse. The resulting magnetisation-transfer ratio (MTR) is sensitive to the proportion of bound proteins, and hence thought to reflect myelin content. Indeed, the MTR is commonly used in the study of demyelination disorders such as multiple sclerosis (Khormi et al., 2023; Zheng et al., 2018). However, the MTR is unlikely to be a selective measure of myelin, because it is also affected by iron concentration, inflammation as well as water content (Mancini et al., 2020; Odrobina et al., 2005; Sled, 2018; Stanisz et al., 2004; Vavasour et al., 2011)(Mancini et al., 2020; Odrobina et al., 2005; Sled, 2018; Stanisz et al., 2004; Vavasour et al., 2011).

Another approach to estimate myelin uses the ratio of signals from T1w and T2w images (Ganzetti et al., 2014; Glasser et al., 2022; Glasser & Van Essen, 2011). This relies on the premise that T1w and T2w contrasts are both sensitive to myelin content, but in the opposite manner, such that their ratio enhances myelin sensitivity (provided appropriate B0 corrections are applied to each image; (Ganzetti et al., 2014; Glasser et al., 2022)). This measure has mainly been applied to study cortical myelin content, but some studies have also used it to examine integrity of WM tracts (Boaventura et al., 2024; Boroshok et al., 2023; Demirtaş et al., 2019; Glasser et al., 2022; Grydeland et al., 2013, 2019; Lee et al., 2024; Pelkmans et al., 2019; Shafee et al., 2015; Sui et al., 2022; Vidal-Piñeiro et al., 2016). However, it is still unclear how the T1/T2 ratio in WM tracts relates to other WM measures. Indeed, some recent work suggests that the T1/T2 ratio in WM is not necessarily sensitive to myelin (Arshad et al., 2017; Sandrone et al., 2023; Uddin et al., 2018).

Most recently, DWI has become the main technique to study WM (Sotiropoulos & Zalesky, 2019). It measures the diffusion of water molecules in vivo, which happens at the micrometric scale during the duration of a typical scan, thereby providing information about microstructural alterations below the scale of the image voxels (Le Bihan & Johansen-Berg, 2012; Moseley, 2002). Indeed, previous work has argued that DWI provides the most sensitive index of WM alterations compared to more conventional structural MRI contrasts such as those described above (Maillard et al., 2013; Nusbaum et al., 2001; Pelletier et al., 2017).

DWI provides multi-dimensional information, to which different mathematical models can be fit in order to extract properties of the diffusion tensor, such as fractional anisotropy (FA) and mean signal diffusion (MSD), and including higher order moments, such as mean signal kurtosis (MSK) (Henriques, Correia, et al., 2021a; Henriques, Jespersen, et al., 2021). These have been indirectly related to the underlying WM microstructure (Basser et al., 1994; Jensen et al., 2005; Jensen & Helpern, 2010). Biophysical microstructural models have also been fit in attempts to estimate parameters with a direct biological interpretation (Huber et al., 2019; Novikov et al., 2018; White et al., 2013). One of the most popular of these is the “Neurite Orientation Dispersion and Density Imaging” (NODDI) model (Zhang et al., 2012), which measures the neurite density index (NDI), fibre orientation dispersion index (ODI) and the free water volume fraction (Fiso). More recently still, Baykara et al. (2016) suggested that the spread of MD values across voxels within WM tracts (a WM skeleton) - which they called the “peak width of skeletonised mean diffusivity” (PSMD) and shown to be particularly sensitive to age-related WM differences. This measure, like WMHI volume, captures WM damage that can, in principle, occur anywhere in the brain within the white matter skeleton.

While many studies have examined the effects of age on at least one of these MRI metrics of WM (e.g., Cox et al., 2016; Davis et al., 2009; Dekkers et al., 2019; Huber et al., 2019; Lawrence et al., 2021; Lebel et al., 2012; Liu et al., 2023; Mendez Colmenares et al., 2023; Schilling et al., 2023; Sexton et al., 2014; Wartolowska & Webb, 2021; Westlye et al., 2010; Yeatman et al., 2014), their simultaneous presence in a large cohort is rare. Furthermore, multiple WM metrics are often tested separately, rather than simultaneously, and their relative sensitivity to ageing and cognition has not been systematically explored. There are prior suggestions that they may be somewhat complementary. For example, Westlye et al. (2010) reported that the macrostructural property of total WM volume (WMV) was not highly correlated with the microstructural property of FA measured from DWI. There are multiple potential reasons for this, such as FA and WMV being differentially sensitive to the presence of non-neuronal cells in WM, and/or the proportion of free water and/or the organisation of axons. Indeed, even within the domain of DWI, Henriques et al. (2023) used factor analysis to conclude that three factors under-pinned their six DWI metrics (FA, MSD, MSK from a tensor model, and NDI, ODI and Fiso from the NODDI microstructural model), indicating both shared but also unique variance explained by their combination.

Here, we started with a similar dimension reduction approach, but now using a total of 11 whole-brain, WM measures: the 6 DWI metrics used by Henriques et al. (2023), plus a PSMD-like measure, together with WMV (T1w), WMHI (T2w), MTR and T1/T2 ratio (see Fig. 1). Apart from examining the relationship of all these measures and whether they can be sufficiently well represented by a small number of factors, we also examined whether the derived factors had convergent construct validity, in terms of being related to other biological and cognitive measures, namely cardiovascular health and cognitive performance.

**Figure 1.**
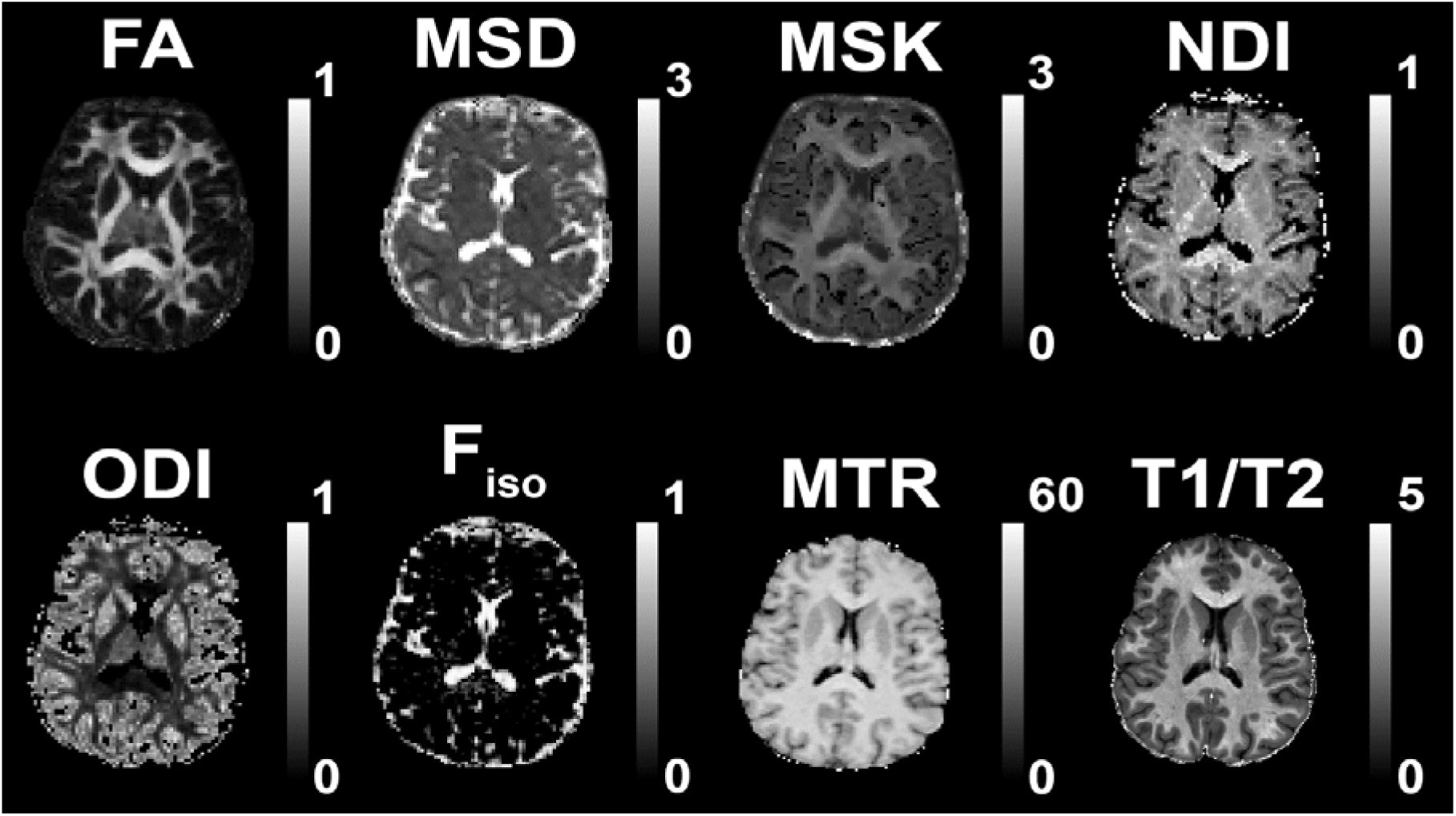
Brain map of the 8 WM measures included in the ROI analyses. We show a young participant’s data from 6 diffusion metrics (FA, MSD, MSK, NDI, ODI, Fiso), plus the additional measures of MTR and T1/T2 ratio. Not shown are the raw T1w+T2w images from which total WM volume (TWM) was calculated, and the other two global measures of White Matter Hyperintensities (WMHI), which were computed across WM ROIs using the T1w and T2w images and the SAMSEG algorithm, and PSMD, which was computed across the WM ROIs using the DWI FA and MSD measures.

The reason for focusing on cardiovascular health was because it has been suggested that WM is particularly vulnerable to hypoxia from reduced cerebral blood flow (Black et al., 2009). This is because the arterioles that supply oxygen to astrocytes and oligodendrocytes are narrow and long, making perfusion more difficult to maintain compared to perfusion in gray matter (Kang et al., 2022; Martinez Sosa & Smith, 2017; Pantoni et al., 1996; Rowbotham & Little, 1965). Indeed, recent work has shown that WM microstructural changes are associated with increased cardiovascular risk in middle and older adults (Kennedy & Raz, 2009; Reas et al., 2021; Wartolowska & Webb, 2022; Wei et al., 2020). Multiple cardiovascular measures have been used in previous studies, including heart-rate and heart-rate variability, body-mass index, static blood pressure, and pulse pressure – the difference between systolic and diastolic blood pressure. Most of these studies have linked a single cardiovascular measure to a single measure of WM (d’Arbeloff et al., 2019, 2021; Daskalopoulou et al., 2017; Graciani et al., 2023; Hofman et al., 2023; Livingston et al., 2020; Moreno-Agostino et al., 2020; Palta et al., 2021; Raichlen et al., 2024; Strömmer et al., 2020). Here, we utilised previous work in the Cam-CAN cohort by King and colleagues (2023, 2024) who showed that common cardiovascular measures can be captured by 3 distinct latent factors. More specifically, we examined how these 3 cardiovascular factors predicted our WM factors.

In terms of the consequences rather than causes of WM health, we examined how our WM factors related to several “fluid” cognitive measures, namely fluid intelligence, processing speed and episodic memory. These cognitive measures were first adjusted for age and sex. To additionally account for other individual differences in cognition in participants, we ran an additional control analysis where we tested the relationship between WM factors and cognition after adjusting for each individual’s polygenic propensity score for high general cognitive ability.

## Methods

### Participants and Materials

We used neuroimaging data from Phase 2 (Arm 1) of the Cambridge Centre for Aging and Neuroscience (Cam-CAN, www.cam-can.org, (Shafto et al., 2014)), which included healthy individuals aged between 18-89, approximately half male/female. None of the participants had a current diagnosis of dementia or mild cognitive impairment, and scored 25 or higher on the mini mental state examination (Folstein et al., 1975). Participants were native English speakers, had no neurological disorders, and had normal or corrected to normal vision and hearing (see Shafto et al., 2014 for further details).

At least some type of valid MR data was available on total of 708 participants. However, for our first question of investigating the number of latent factors (and relative importance of different WM measures (see Fig. 1), including whether non-DWI sequences add complementary information), we used only participants who had data on all of the MR sequences, to avoid biasing the decomposition method towards the T1w and T2w sequences (which were the most common across participants). A total of 583 participants had valid data for all four MR sequences (T1w, T2w, MTw, DWI). We subsequently removed 13 participants who had outlier scores on at least one of the 11 WM measures described below, leading to a final sample of n=570 (see Fig. 2). Outliers were defined as having residual values more than 5 SDs from the mean, after accounting for age and sex effects.

**Figure 2.**
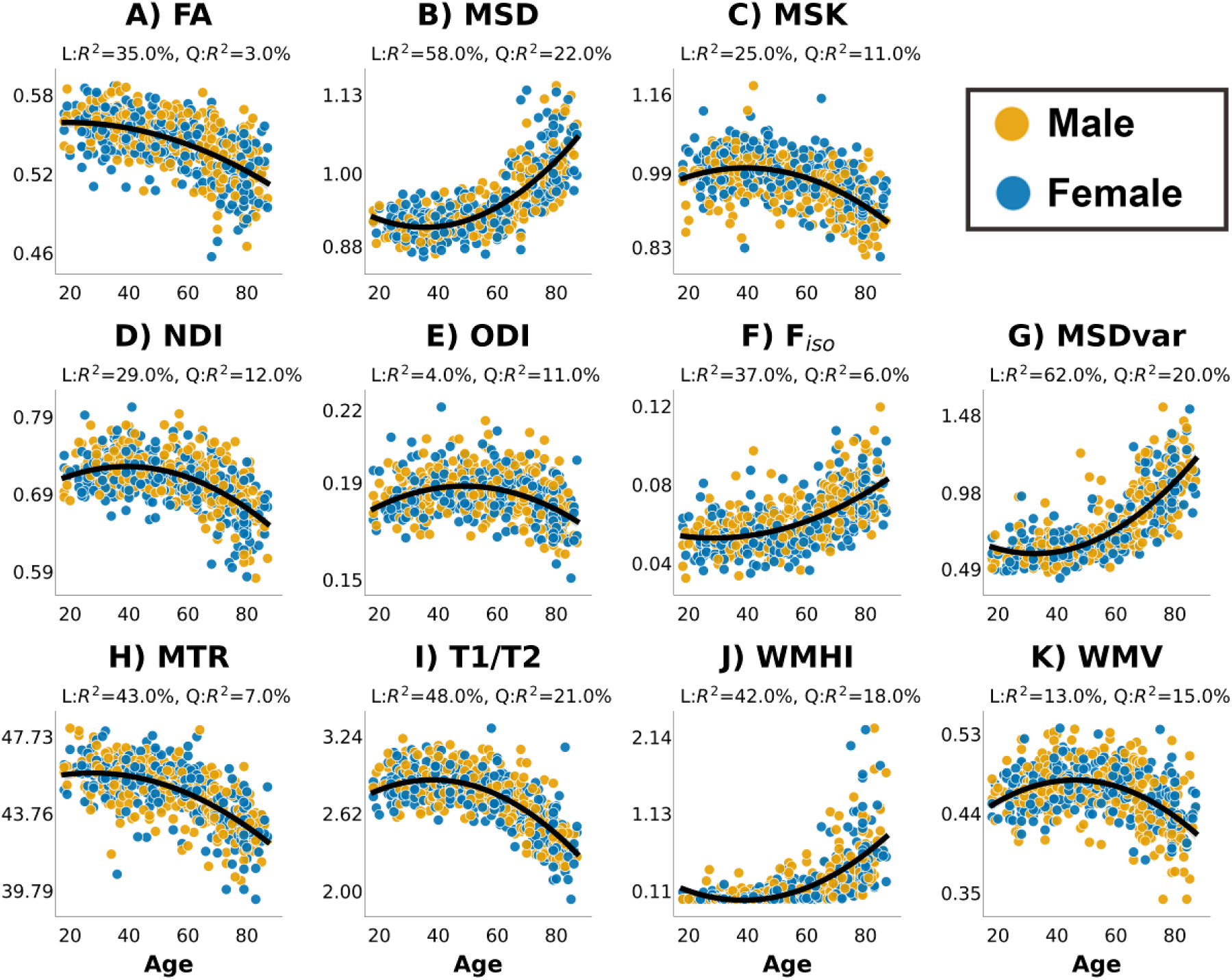
Values for all 11 WM metrics by sex and across age after removing outliers. A) Fractional Anisotropy (FA); B) Mean Signal Diffusion (MSD); C) Mean Signal Kurtosis (MSK); D) Neurite Density Index (NDI) from NODDI; E) Orientation Dispersion Index (ODI) from NODDI; F) Volume Fraction of Free isotropic water (Fiso) from NODDI; G) MSDvar - range of MSD values across 27 WM tracks; H) MTR values across age; I) T1w vs T2w ratio; J) Volume of White Matter Hyper-Intensities (WMHI) normalised by head size. K) total White Matter Volume (WMV) volume corrected for head size. The proportion of variance (R^2^) explained by the Linear (L) and Quadratic (Q) effects of age (after accounting for main effect of sex and sex-by-age interactions) is shown at the top of each panel.

For the second question, where we related WM factors to cardiovascular health and to cognition, we maximised our sample size by including all N=708 participants for whom we had any WM, cardiovascular, or cognitive measure, and used Full-Information Maximum Likelihood (FiML) methods to impute missing data (Graham, 2003; Little & Rubin, 2019; Rosseel, 2012). Outlier values were also treated as missing data. For completeness, there were n = 607 participants with DWI measures; n = 584 with MTR values; and n = 640 with metrics derived from T1w and T2w sequences (TWM, WMHI and T1/T2).

### Cardiovascular Measures

For cardiovascular measures, we used blood pressure (BP, n = 579), pulse pressure (PP, n = 579), heart rate (HR, n = 580), heart rate variability after low- and high-pass filtering (HRV_LF/HF, n = 604) and body-mass index (BMI, n = 587).

### Cognitive Measures

For cognitive measures, we chose the fluid abilities of fluid intelligence (n=660), processing speed (n = 658) and episodic memory (n=706). Fluid intelligence was measured using the Cattell Culture Fair test, which included 4 sub tests (Cattell, 1971). We performed factor analysis to reduce the four sub-tests into one factor. Processing speed was captured across two tasks - simple and choice reaction time (RT) tasks – both of which involved one of four lights triggering one of four finger presses (full details in Shafto et al., 2014). The mean and standard deviation of RTs across 40-60 correct trials were computed for each task. To account for the fact that both mean and standard deviation of RTs were positively skewed, we inverted RTs before computing mean and standard deviation (Kievit et al., 2016; Ratcliff, 1993; Tsvetanov et al., 2016). These four measures were then standardised, and reduced to a single latent variable using factor analysis. Episodic memory was measured using immediate recall, delayed recall and delayed recognition memory scores from the Wechsler Logical Memory task (Wechsler, 1991), and factor analysis was again used to reduce these three measures into a single factor. For all factor analyses, missing data were imputed Full Information Maximum Likelihood.

### MRI Acquisition

All imaging data were collected on a Siemens Trio 3T MRI scanner with 32-channel head coil. T1w and T2w images had 1mm isotropic voxel size, FOV = 256 x 240 x 192 mm and flip angle=9°. The T1w was acquired with an MPRAGE sequence with TE of 2.99ms and TR of 2250ms; the T2w was acquired with a SPACE sequence with TE of 408ms and TR of 2800ms. Both were acquired with acceleration factor 2 using GRAPPA sequence.

MTw images were collected with voxel size of 1.6mm isotropic, FOV = 192 x 192 x192 mm, TE = 5ms, and flip angle = 12°, acquired using a spoiled gradient echo sequence with TR = 30ms or TR = 50ms, depending on whether the participant’s specific absorption rate (SAR) estimation for the TR = 30ms sequence exceeded the stimulation limits. Two images were acquired, with or without a Gaussian RF saturation pulse with an offset frequency of 1950 Hz (bandwidth = 375 Hz, 500° flip angle, duration = 9984 µs).

DW images were collected with voxel size of 2mm isotropic, FOV = 192 x 192 x 132 mm, TE = 104ms, TR = 9100ms, acquired using echo-planar imaging with twice refocused spin echo (TRSE) to reduce eddy-current artefacts. Data were collected with two non-zero b-values (1000 and 2000 s/mm^2^) along 30 diffusion gradient directions and for three b = 0 volumes, with partial Fourier of 7/8, and acceleration factor of 2 using GRAPPA with 36 reference lines. More information about the MRI acquisitions is reported in Taylor et al. (2017), while full sequence parameters are here: https://camcan-archive.mrc-cbu.cam.ac.uk/dataaccess/pdfs/CAMCAN700_MR_params.pdf.

### Image pre-processing

The T1w and T2w images were initially processed used the SPM12 software (Wellcome Centre for Human Neuroimaging; https://www.fil.ion.ucl.ac.uk/spm), release 4537, implemented in the Automatic Analysis (AA) pipeline, release 4.2 (Cusack et al., 2015). Pre-processing is described in Taylor et al. (2017), but in brief, the T2w image was rigid-body coregistered to the T1w image, and then both were rigid-body registered to a Montreal Neurological Institute (MNI) template brain (just to improve starting point for segmentation). Both images were then corrected for B0 bias, after which they were combined for purposes of segmentation using SPM’s multimodal, unified segmentation and normalisation (Ashburner & Friston, 2005). We computed total white matter volume based on the native space segmentation maps. Total inter-cranial volume (TIV) was used to normalise the volumetric white matter measures.

As part of the AA pre-processing, we used the DARTEL algorithm (Ashburner, 2007) to create a mean gray-matter template across the whole sample of participants, which was subsequently 12-parameter affine transformed to MNI space. The inverse of these combined transformations was then applied to convert the JHU ROIs back into the native space image of each participant’s T1/T2 and MTR images (see next section).

The raw T1w and T2w images were also processed with the MRTool within SPM12 (Ganzetti et al., 2014). The T2w image was coregistered to the T1w image using a rigid-body transformation. A transit field intensity inhomogeneity (B1 field) correction was then applied to each image separately, since they might have different intensity non-uniformity. A new image was then created by dividing the T1w value by the T2w value at each voxel.

The two MTw weighted images were rigid-body coregistered and used to calculate the magnetisation transfer ratio (MTR) as (M0 - Ms)/M0, where Ms is the mean signal intensity with the saturation pulse and M0 is that without saturation.

The DWI data were pre-processed with a common pipeline that has been described in detail in Winzeck (2021). DIPY (https://dipy.org/) procedures were implemented to 1) reduce noise via PCA (Veraart et al., 2016) and 2) correct Gibbs artefacts using an adapted sub-voxel shift procedure (Henriques, 2018; Kellner et al., 2016) Eddy current correction was not applied since the eddy artefacts were minimised by the TRSE sequence (Reese et al., 2003).

Six DWI metrics for each voxel were computed as described in Henriques et al. (2023): (1) Fractional Anisotropy (FA) computed from standard diffusion tensor imaging (DTI); (2) Mean Signal Diffusion (MSD) and (3) Mean Signal Kurtosis (MSK) from the Mean Signal Diffusional Kurtosis Imaging (DKI) (Henriques, Correia, et al., 2021b); and (4) Neurite Density Index (NDI), (5) Orientation Dispersion Index (ODI), and (6) isotropic Free water volume fraction (Fiso) estimated from Neurite Orientation Dispersion and Density Imaging (NODDI) modelling. Some illustrative images of these 6 diffusion metrics, and the MTR and T1/T2 estimates, are shown in Figure 1.

### Region of Interest (ROI) definition

WM tract ROIs were defined by the John Hopkins University (JHU) atlas (Mori et al., 2008). WM metrics were obtained by first averaging across all voxels within the 48 WM tracts, and then averaged again over homotopic ROIs to create 27 WM ROIs. For the DWI data, the ROIs were back-projected from an FA template into each person’s FA native space using FSL’s linear and non-linear registration tools (Jenkinson et al., 2012; Smith et al., 2004; Woolrich et al., 2009). To suppress the impact of free-water partial volume effects from cerebral spinal fluid (CSF), and to minimise the impact of degenerative ODI and NDI estimates in voxels containing mostly free water, voxels with Fiso values larger than 0.9 were removed from the ROIs. For the T1/T2 and MTR data, we used the same ROIs that were inverted back to native space using the inverse DARTEL warps derived as a part of the AA pipeline described above.

To match the whole-brain metrics below, the WM metrics were averaged across the 27 ROI tracts to create a single global measure of white matter integrity from each measure.

### Whole-Brain Metrics

We extracted total white matter (WMV) volume from SPM 12’s segmentation procedure. We then regressed out total intracranial volume (TIV), also estimated by SPM12, to ensure the measure was not confounded by head size.

While WMHIs are usually measured by a FLAIR (Fluid-Attenuated Inversion Recovery) sequence, recent developments of the SAMSEG algorithm allow them to also be estimated from T1w and T2w volumes (Cerri et al., 2023). The SAMSEG algorithm is a multi-atlas probabilistic segmentation method that has been shown to be robust to noise and the inherent intensity inhomogeneity in the MRI data. For each voxel, the SAMSEG algorithm outputs a probability that it contains a white matter hyper-intensity lesion. We used a probability threshold of 0.5 to define binary lesion masks across the whole brain, from which we computed WMHI lesion volumes. We then normalized this WMHI volume by TIV, like for WMV above.

The last measure represented the range of values across voxels present in the mean signal diffusion (MSD) image, and is conceptually identical to the recently proposed measure of “peak width of skeletonised mean diffusivity” (PSMD) (Baykara et al., 2016). The main difference here was that we computed the 90% percentile range of MSD values for all voxels within the 27 ROI tracts, rather than within an FA skeleton. This “MSDvar” measure was highly correlated with the standard PSMD measure (r = 0.86) taken from King et al. (2024).

### Statistical Analysis

We had two main aims in this manuscript. The first one was to summarise the relationship between different WM measures. Specifically, we wanted to examine which WM measures are strongly interrelated with others and whether non-DWI measures provide complementary information not captured by DWI metrics. The second main question was to examine how WM health relates to cognition and Cardiovascular measures. The derived white matter measures are available as csv files hosted on osf. We also provide R code to reproduce the main analyses https://osf.io/y7ct8/. Below we describe the approaches for the 2 main questions in the paper.

#### 1. Using Principal Component Analysis to test relationships between WM measures

Principal Component analysis (PCA) was applied to all 11 WM measures, after Z-scoring each measure separately. We used recently developed, element-wise cross-validation to identify the number of components, as implemented in the MEDA Matlab toolbox (Camacho et al., 2015; Saccenti & Camacho, 2015b). This element-wise k-fold cross-validation (*ekf*) has been shown to be more appropriate than the standard whole observation (holding out whole row) cross-validation for PCA (Bro et al., 2008; Camacho & Ferrer, 2012; Josse & Husson, 2012; Saccenti & Camacho, 2015a, 2015b). Note however that ekf is only one of many ways to estimate number of components, and each method seems to be differentially sensitive to different aspects to the data (Saccenti & Camacho, 2015a). After determining the number of components, we used the “Varimax” rotation to enhance interpretability.

#### 2. Using path models to validate WM factors against cardiovascular health and cognition

Here we attempted to validate the above WM factors by relating them to independent physiological and cognitive data. We ran two separate path analyses, using the lavaan R package (Rosseel, 2012). In one path model, we examined how the WM factors were predicted by latent cardiovascular factors, in line with the idea that cardiovascular problems lead to white matter damage. In the second path model, we examined how the WM factors predicted the cognitive measures of fluid intelligence, processing speed and episodic memory. In other words, the two path models allowed us to examine both the potential causes (cardiovascular) and consequences (cognitive) of the WM factors. In both of these path models we show paths with and without controlling for linear and quadratic age effects and sex.

Note that some cohorts may not have, or be able to acquire, all the WM metrics utilised here. Therefore in Supplementary materials, we also report how each individual WM measure was independently related to cognition, so that researchers might be able to choose the ‘best’ metric (and MR sequence) if their goal is to relate WM to cognition (see Supplementary Figures 3 & 4).

##### Covariates

We report the two path models both with and without including age effects on the outcome variable in the same model. Regressing out age effects can remove some of the variance in WM factors caused by age-related but non-cardiovascular variables, and some of the variance in cognition caused by age-related but non-WM variables. However, age is also likely to cause individual differences in cardiovascular and/or WM, so the danger of regressing it is that true cardiovascular-WM and/or WM-cognition relationships are masked. This is a limitation of cross-sectional data (Deary et al., 2009; Raz & Lindenberger, 2011; Walhovd et al., 2023), but such data can at least be used to identify potential links between these variables that can be further tested in longitudinal data.

Another source of individual differences in brain and cognition is genetics. A large proportion of the between-person variance in intelligence is attributable to early-life differences in intelligence (Deary et al., 2010). Given that intelligence is highly heritable (Akimova et al., 2021), we also adjusted for potential genetic effects in the second path model in which WM predicted cognition. Prior work in Cam-CAN has exampled single gene effects (Henson et al., 2020; P. P. Raykov et al., 2025), but here we used a polygenic risk score (PRS) for general cognitive ability with the single nucleotide polymorphisms (SNP) effect sizes identified in a GWAS study from other independent cohorts (Savage et al., 2018). The PRS was computed, independently of our other cognitive and brain measures, using the Bayesian continuous shrinkage prior approach, which robustly infers the posterior effect sizes of SNPs, while also taking into account the external linkage disequilibrium (Ge et al., 2019). PRS was adjusted for the first 10 genetic principal components (to control for population stratification effects), and then standardised.

## Results

We first explored each of the 11 WM measures in terms of the relationship with the age and sex of each participant (Figure 2). The results of the full regression models are available online (see Methods), so just summarised here.

Most measures exhibited a main effect of sex, except for MSD, ODI, MTR, WMHI and WMV. The measure with largest sex difference was MSK, where females tended to have higher values (β = 0.007, t = 3.80 p < 0.001, η^2^ = 0.025), though this effect size was still relatively small.

All measures showed a large, main effect of age (see Figure 2 for effect sizes for Linear and Quadratic components). The results for the 6 DWI measures were qualitatively very similar to those reported by Henriques et al. (2023), so we focus on the 5 new measures. The “MSDvar” DWI metric tended to increase quadratically with age, similar to the mean MSD values. Indeed, both of these MSD metrics showed the strongest effect of age (with linear and quadratic components combining to explain 79% and 82% of their variance respectively). On the other hand, MTR and T1/T2 ratio decreased quadratically with age. There was a strong non-linear increase in WMHIs with age, which were most prominent in old age, in line with prior work (d’Arbeloff et al., 2019; Lyall et al., 2020; Moon et al., 2018; Raichlen et al., 2024; Ritchie et al., 2024; Torres et al., 2015; Vergoossen et al., 2021; Wartolowska & Webb, 2021). WMV exhibited an inverted U-shape relationship across the whole age range.

### Principal Component Analysis for the 11 WM metrics

Figure 3 shows the correlation among the 11 WM measures, before (left) and after (right) adjusting for second-order effects of age, sex and their interactions. We first regressed out the confound variables and then computed correlation of the residuals. We note this is more aggressive clean-up as likely there is variance shared across white matter measures and age. The high similarity between some measures even after partialing effects of age indicates that the relationships between metrics are not driven solely by common age dependency. In particular, high positive correlations are observed among Fiso, WMHI, MSD and MSDvar, and between FA, WMV, MTR, T1/T2, MSK and NDI.

**Figure 3.**
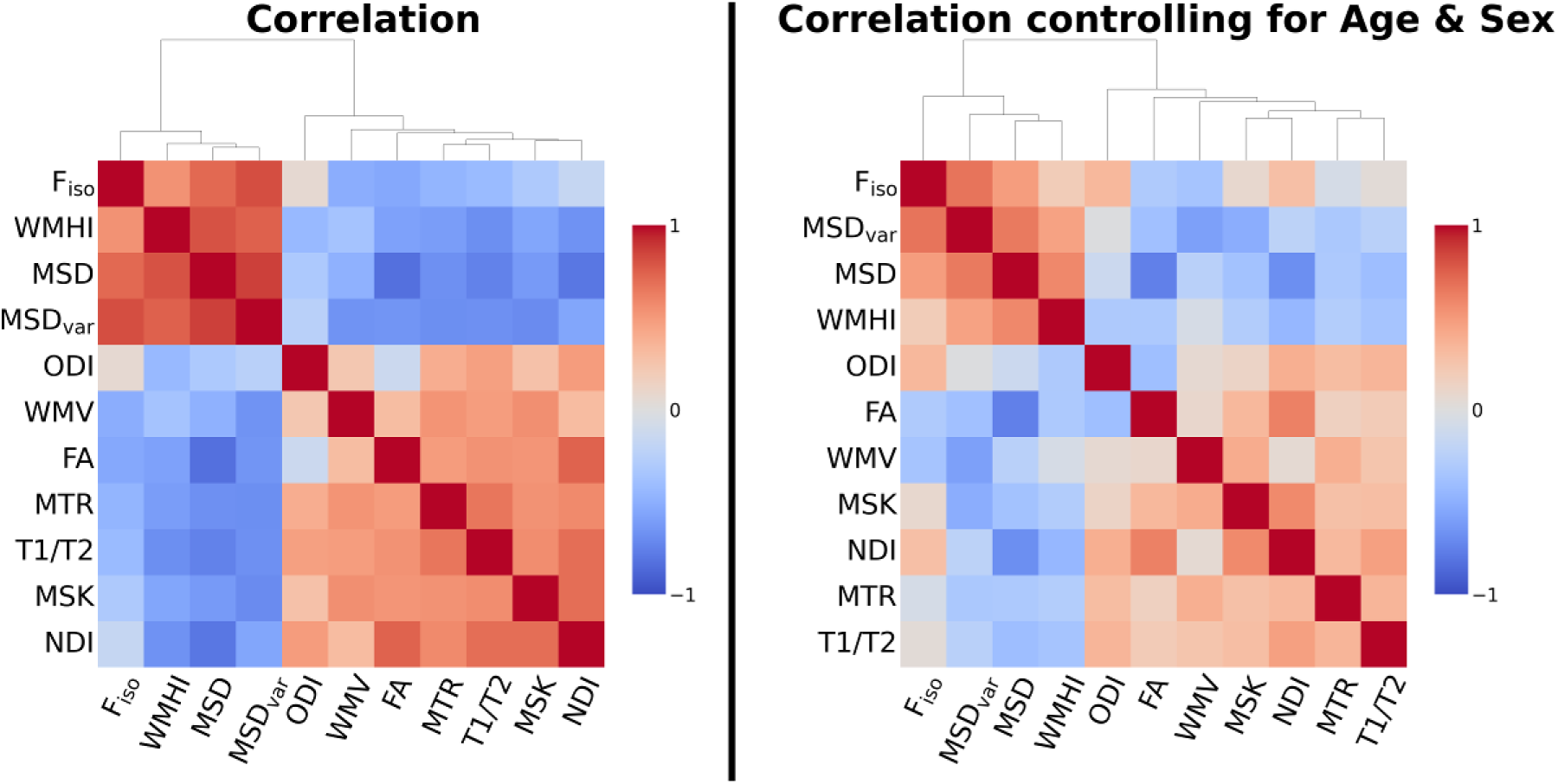
Correlation matrix for all 11 WM measures across participants before (left) and after (right) correcting for sex, linear and quadratic age effects, and the sex-by-age interactions. Metrics are ordered according to the dendrogram of agglomerative similarity between measures.

We used ekf cross-validation to determine the number of PCA components. We note that his element-wise cross-validation methods has recently been suggested as a robust way to determine number of PCA components (Camacho & Ferrer, 2012; Saccenti & Camacho, 2015b). To ensure our factors were interpretable we performed Varimax rotation on the 4 components.

The PCA with 4 components captured 88.7% of the variance in the 11 WM measures. The 1st principal component captured 59.2% of the variance, the 2^nd^ PC explained 13.4%, the 3^rd^ explained 9.9% and the 4^th^ component explained 6.3% of the variance. The 5^th^ component that was not retained captured 3.7% of the variance (Figure 4).

**Figure 4.**
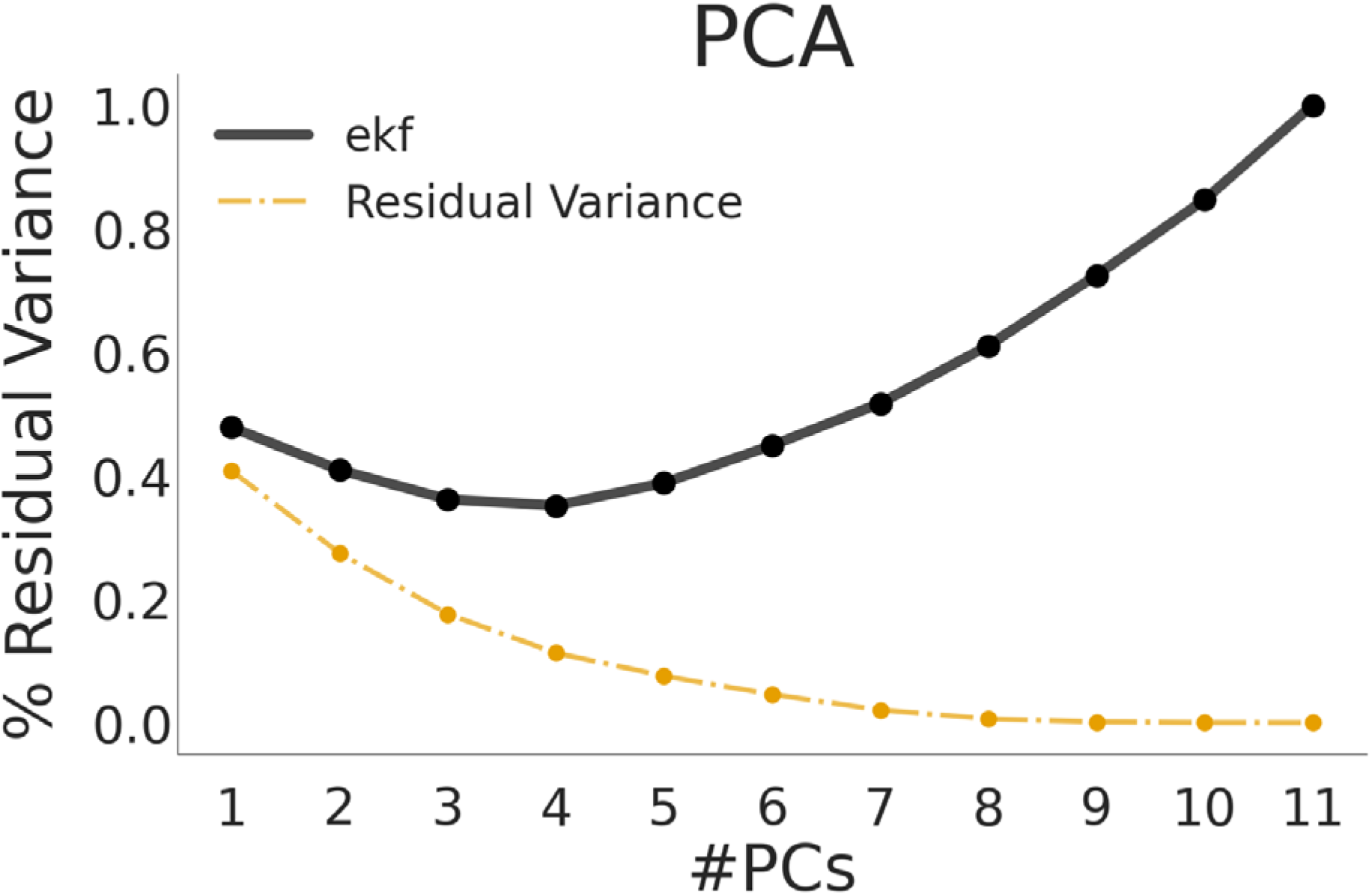
PCA cross-validation. The figure shows the ekf cross-validated error as a function of number of principal components retained, along with the residual variance (inverse of variance explained) in orange. Based on the ekf, we selected 4 PCs and performed factor analysis with 4 factors.

Figure 5 shows the loadings for each factor and how the factor scores related to age. Overall, Factors 1, 2 and 3 resembled the factors found in Henriques et al. (2023). The first factor loaded strongly on FA and NDI, with a negative loading on MSD. This factor declined with age and likely reflects microstructural properties such as axonal fibre density and myelination. Interestingly, Henriques et al. also found that MSK loaded on this factor, whereas here it loaded more on the additional, fourth factor (see below). Factor 1 also loaded positively on MTR and T1/T2 ratio, consistent with age-related reductions in myelin content, and negatively on WMHI, consistent with demyelination with age.

**Figure 5.**
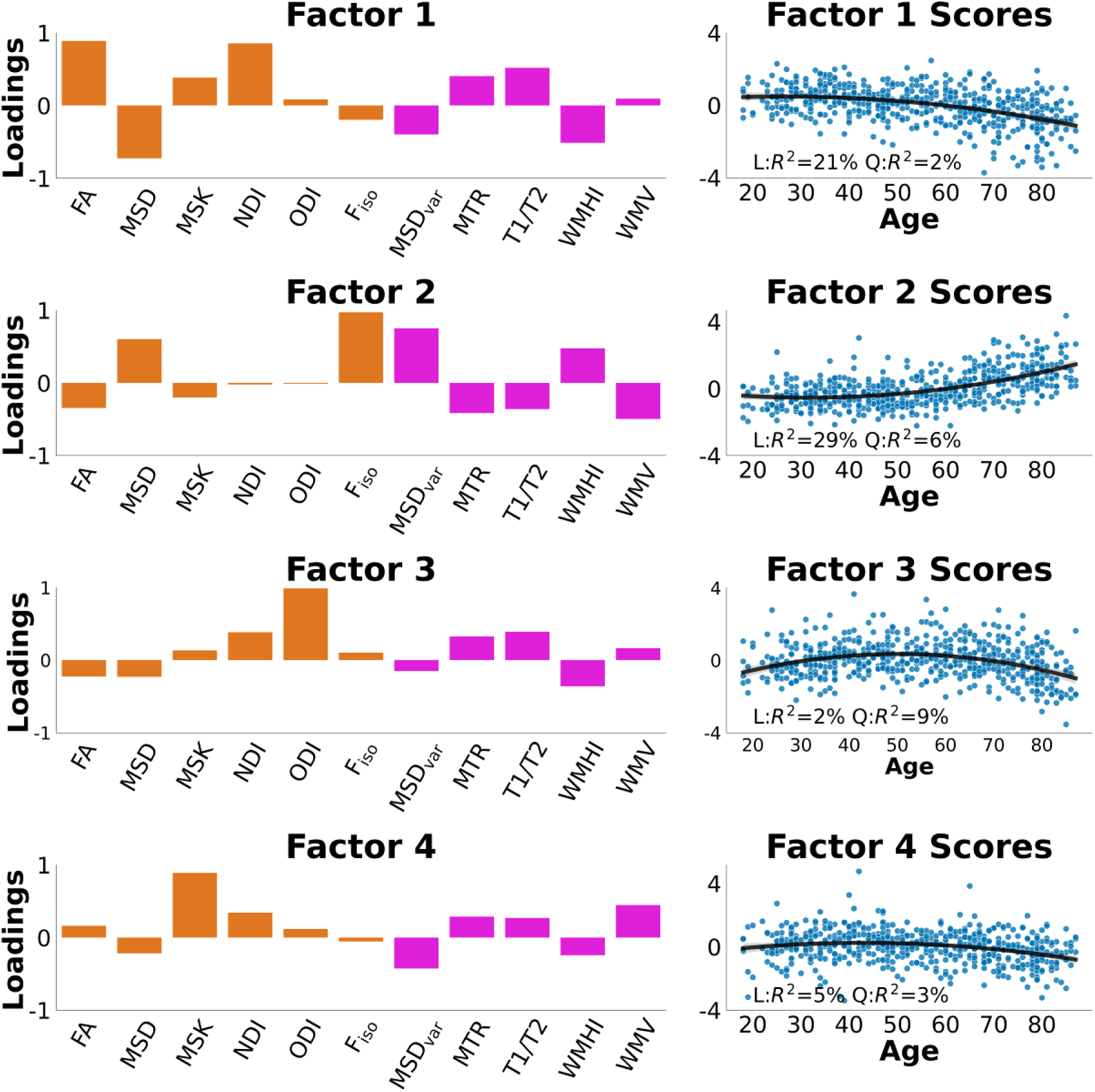
Loadings and Factor scores across age. We show factor loadings across all 11 WM measures and how the factor scores varied with age. We interpret Factor 1 as a microstructural properties/myelination factor. Factor 2 as free-water/tissue damage factor. Factor 3 as fibre-crossing complexity factor. Factor 4 as microstructural complexity

Factor 2 loaded positively mainly on Fiso and MSD, most likely representing increases in free water content. This factor increased with age. Interestingly, MSDvar – the DWI measure conceptually identical to PSMD - also loaded positively on this factor, as did WMHI. Conversely, MTR, T1/T2 ratio and WMV all loaded negatively on this factor to some extent. This suggests that Factor 2 captures WM damage/lesions leading to increase partial volume effects between tissue and CSF.

Factor 3 mainly loaded positively on the ODI metric, with weak loadings on all other metrics, and showed a weak, inverted-U shape function of age. Like in Henriques et al. (2023), this factor seems to relate to complexity of fibre configurations (such as crossing fibres and fibre dispersion).

Factor 4, which was not needed in Henriques et al.’s analysis of DWI metrics alone, loaded positively on MSK, with a smaller positive contribution from WMV and a smaller negative contribution from MSDvar. It showed a weak decline with age. Interpretation of this factor is less clear but appears to reflect general microstructural complexity / volume that is distinct from fibre density/myelination and free-water/lesions (see also Kamiya et al., 2020).

### Validating the white matter factors against cardiovascular health and cognition

#### Cardiovascular measures predicting white matter

First, we examined whether the latent vascular scores predicted the latent WM scores. We defined 3 cardiovascular factors based on prior work from the Cam-CAN cohort (King et al., 2023). The first latent vascular factor (LVF1) captured static blood pressure and related to BMI; the second (LVF2) captured pulse pressure and heart rate; the third (LVF3) loaded highly on both high and low frequency heart rate variability. We tested how these cardiovascular factors predicted the four WM factors (Supplementary Fig 1). We had data from 627 participants.

We first report the path model relating cardiovascular and WM factors (WMFs) without adjusting the WMFs for age and sex effects (Figure 6, top left; see Supplementary Table 1 for full parameters (Thériault, 2023)). This model would be appropriate if the effects of age and sex on WM were via their effects on cardiovascular health. This model showed WMF1 (reflecting fibre density/myelination) was negatively related to LVF2 (largely pulse pressure) but positively related to LVF3 (heart-rate variability). This is consistent with low pulse pressure and high heart-rate variability being associated with better cardiovascular health.

**Figure 6.**
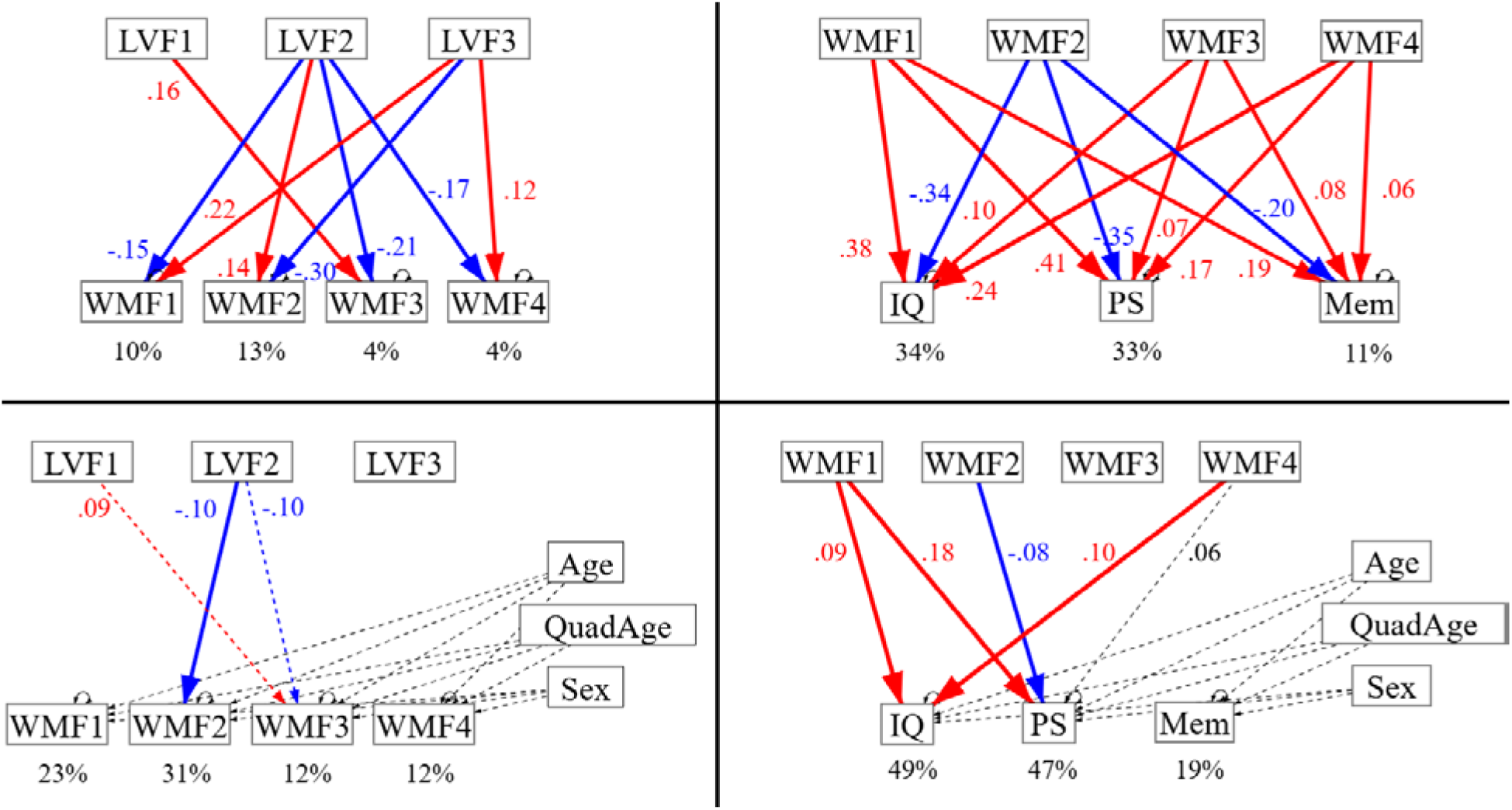
Path models relating cardiovascular factors to WM factors (left panels) and WM factors to Cognitive factors (right panels), without (top panels) and with (bottom panels) adjustment of outcome variables for Age and Sex effects. LVF = latent vascular factor; WMF = white matter factor; IQ = fluid intelligence; PS = processing speed; Mem = episodic memory.

WMF2 (reflecting free-water content / WM damage) showed the opposite pattern of a negative path from LVF2 and positive path from LVF3, this time generally consistent with poor vascular health being associated with more WM damage. WMF3 (reflecting fibre-configuration complexity) was positively related to LVF1 (largely static blood pressure) and negatively related to LVF2 (pulse pressure). The reason for these is less clear. Finally, WMF4 was negatively related to LVF2 and positively related to LVF2, like WMF1.

We also report results from a model where (linear and quadratic) effects of age and sex affected WM through routes that are independent of cardiovascular heath (Figure 6, bottom left). Because age in particular has such strong effects on WM, the only path from cardiovascular factors that remained significant was the path from LVF2 to WMF2, though this was now negative rather than positive (see Discussion). The paths to WMF3 from LVF1 and LVF2 also approached significance (Supplementary Table 2).

### White Matter predicting Cognition

The cognitive factors of fluid intelligence (IQ), processing speed (PS) and episodic memory (Mem) were based on theoretical distinctions rather than data-driven factor analysis (though each factor was itself estimated from factor analysis of several component scores; see Methods). Without accounting for age and sex, all WM factors were related to all cognitive factors (Figure 6, top right; Supplementary Table 3). As expected, WMF2 (reflecting free-water content / WM damage) was negatively related to all three cognitive factors, whereas the remaining three WM factors were positively related to all cognitive factors, consistent with good WM health being associated with better cognitive outcomes. Nonetheless, these relationships could all be driven by independent effects of age on both WM and Cognition.

In the path model that allowed for such independent effects of age (and sex) (Figure 6, bottom right and Supplementary Table 4), the positive effects of WMF1 on fluid intelligence and processing speed remained significant. The positive effect of WMF4 on fluid intelligence also remained significant. Finally, the negative effect of WMF2 on processing speed remained significant. Thus at least some contributions of WM to cognitive abilities appear independent of age, at least for fluid intelligence and processing speed.

### White Matter predicting Cognition Controlling for polygenic effects

Finally, Cam-CAN cohort also has a polygenic score (PRS) for general cognitive ability. We therefore ran a final path model in which the cognitive outcome variables were also adjusted for this PRS. As expected, this PRS was positively associated with all our cognitive measures (Supplementary Table 5). This score is only available in a sub-sample of n= 565 participants, but due to missingness in both neuroimaging and cognitive data, the final path model included only 505 participants.

The positive paths between WMF1 and WMF4 and fluid intelligence and processing speed remained significant even when controlling for this genetic contribution (Figure 7 and Supplementary Table 5), though the negative path from WMF2 to processing speed no longer reached significance (though this could also reflect the lower power from this reduced subset of participants). Interestingly, the PRS was not significantly correlated with any of the WM factors (Supplementary Table 5), consistent with the associations between WM and cognition being at least partly due to environmental and lifestyle factors.

**Figure 7.**
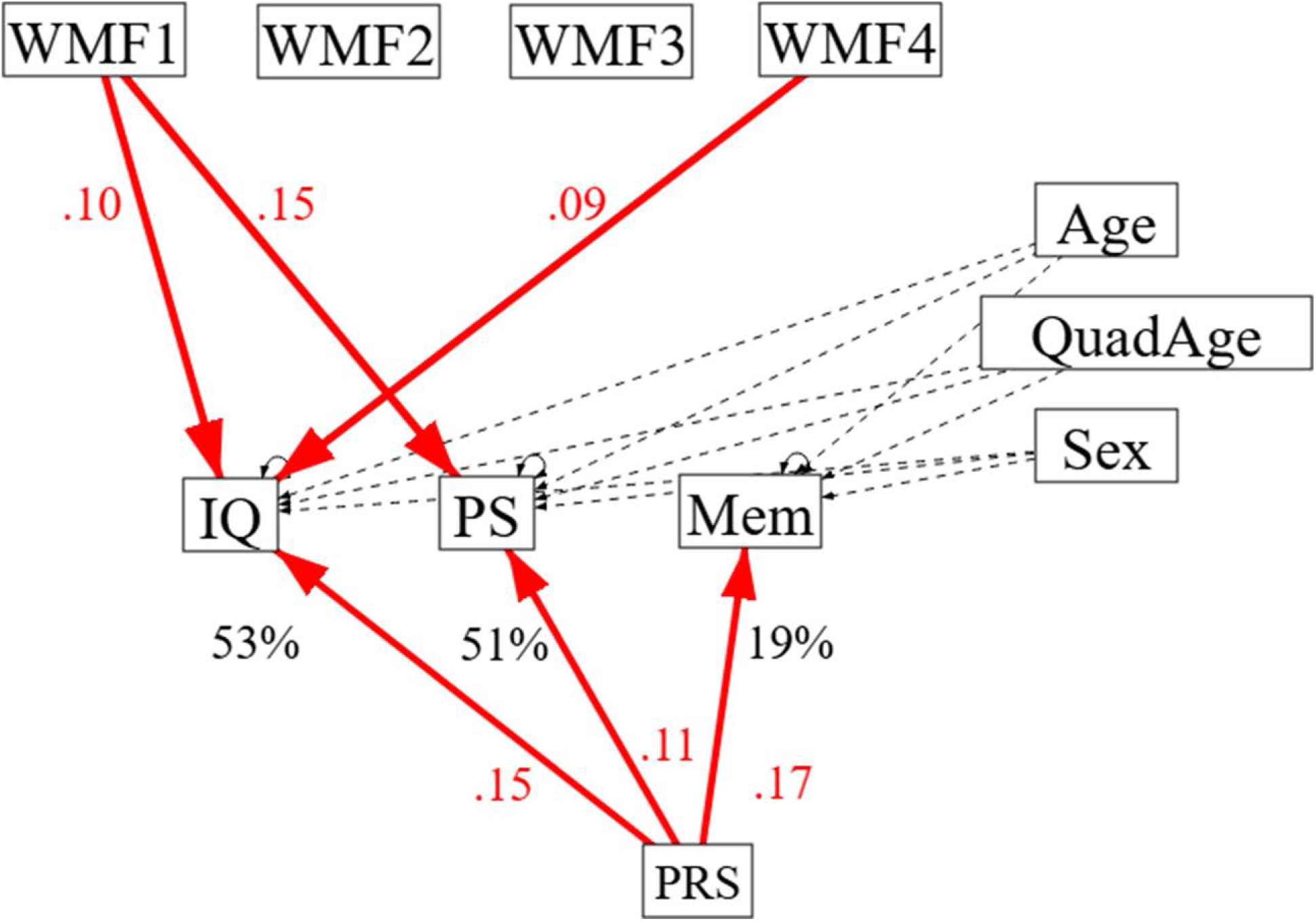
Path Model for effects of WM on Cognition, now additionally controlling for polygenic scores (PRS) for cognitive ability.

## Discussion

Using a large sample of healthy adults, we examined the relationship between 11 different white matter (WM) metrics, derived from 4 different MR contrasts / sequences (T1w, T2w, MTR and DWI), their relationships with age and sex, their underlying factor structure, and the relationship of those factors with cardiovascular health and cognitive abilities. Cross-validated PCA suggested that there are 4 WM factors, which captured 89% of the variance in the 11 metrics. After varimax rotation, these factors seemed to represent fibre density / myelination (WMF1), free water / WM damage (WMF2), crossing-fibre complexity (WMF3) and general microstructural complexity / volume (WMF4). Path models showed that many of these factors were affected by 3 latent factors derived from 6 measures of cardiovascular health, at least one of which (the negative effect on WMF2 of a vascular factor related to pulse pressure) was independent of any shared effects of age or sex. Moreover, many of these factors predicted the cognitive abilities of fluid intelligence, processing speed and episodic memory, even after adjusting for age and sex, at least in the case of fluid intelligence and processing speed. Indeed, the positive effects of WMF1 and WMF4 on fluid intelligence remained even after further adjusting for polygenic disposition for high cognitive ability, consistent with environment/lifestyle contributions.

Our PCA results highlight that multiple potential WM properties underlie common MR contrasts, consistent with previous studies (e.g., Chamberland et al., 2019; Henriques et al., 2023). This suggests that a single MR metric is insufficient to capture fully the effects of variables like age and cardiovascular health on WM. Our previous study identified 3 components underlying 6 metrics from diffusion-weighted MRI (Henriques et al., 2023). Here, the addition of a further 5 metrics, 4 of which were from 3 new MR contrasts, resulted in the need for a fourth component, suggesting some complementary information in T1w, T2w and MTR sequences.

However, the fourth component only explained an additional 6.3% of variance in the 11 WM metrics, and other methods for identifying the optimal number of principal components may suggest fewer than four (there is no universally agreed method for such dimension reduction). Indeed, if three components are in fact sufficient, then one could argue that DWI data alone are sufficient to capture of all these (when used to derive multiple diffusion-related measures, as here), and that there is no additional benefit of acquiring T1w, T2w and MTR data (Supplementary Table 8). In fact, the first three factors here seemed to correspond largely to the three metrics derived from the NODDI model of DWI data.

Nonetheless, even if three components are sufficient, the addition of the 5 extra WM metrics in the present paper helped bolster the interpretation of the WM factors that resulted. Furthermore, extracting latent factors from more WM metrics should in theory improve the accuracy (signal-to-noise ratio) of those factor loadings, by virtue of greater averaging over independent noise sources associated with each metric. Conversely, if one had to choose just one WM metric (from one MR contrast), then Supplementary Figure 3 and Supp. Table 6 shows that MSDvar is likely to be most sensitive to age, MSK is likely to be most sensitive to fluid intelligence, FA is likely to be most sensitive to processing speed, and MTR is likely to be most sensitive to episodic memory (adjusting for age in the latter three cases).

We only considered “global” measures, averaged across 27 ROIs of the JHU atlas. This was because some of the common WM metrics that we wanted to include, such as PSMD are by their nature only defined across multiple WM regions/voxels (here, defined in terms of the 95-5% range of MSD values across ROIs). It is possible that there is regional variation in the underlying WM factors (WMFs), which could even increase the number of components needed (see Henriques et al, 2023, for PCA across ROIs too). Indeed, certain WM metrics might be better suited for certain parts of the brain. This could be the topic of future research.

We sought external validation of the four WM factors by relating them to potential cardiovascular causes. More specifically, we identified 3 vascular factors (LVFs) from 6 measures in CamCAN (systolic and diastolic blood pressure, body-mass index, heart rate, and heart-rate variability after low- and high-pass filtering of ECG), based on prior work by (King et al., 2023). Significant relationships were found between most of the LVFs and most of the WMFs, providing convergent construct validity for the WMFs.

When we adjusted the WMFs for linear and quadratic age effects, many of the contributions from LVFs were no longer significant. With cross-sectional data as here, it is difficult to determine whether the observed relationships between LVFs and WMFs are simply artefacts of independent correlations with age. Alternatively, the reduction in this relationship after adjusting for age can reflect the fact that age is the primary driver of vascular health (Lakatta & Levy, 2003) with age-related variation in LVF subsequently causing variation in WM. It is likely that both of these scenarios are true to some extent. Nonetheless, at least one path survived adjustment for age: that from the LVF representing pulse pressure and the WMF capturing free water / WM damage. This is in line with prior work showing a relationship specifically between pulse pressure and WM health in middle-aged and older adults (Kennedy & Raz, 2009; King et al., 2024; Reas et al., 2021; Wartolowska & Webb, 2022; Wei et al., 2020). Note however that the path was negative, such that higher pulse pressure appeared protective. This was actually a flip in sign compared to when age was not adjusted for (when the relationship was positive, as expected). This suggests that age is a common cause of both increased pulse pressure and WM damage. We note that similar reversal of direction of effect when age is included was observed in previous cross-sectional work in the UK Biobank (Wartolowska & Webb, 2022), highlighting the need for longitudinal and treatment studies to better understand the causal mechanisms through which pulse pressure is associated with WM damage.

Further convergent evidence for the validity of the WMFs was obtained when relating them to cognition. Even after adjusting for age and sex effects, we found that the WMF associated with fibre complexity / myelination and WMF associated with microstructural complexity / volume predicted unique variance in fluid intelligence. The WMF associated with fibre complexity / myelination also predicted processing speed, but this time there was also a unique (and negative) contribution from the WMF associated with free water / WM damage. The fact that the WMF associated with fibre complexity / myelination loaded on FA and NDI and MSD measures is consistent with previous studies that have found relationships between these individual metrics and fluid intelligence and/or processing speed in middle aged and older cohorts (Corley et al., 2023; Cox et al., 2016; Cox, Ritchie, et al., 2019; Grieve et al., 2007; Hernández et al., 2013; Jacobs et al., 2013; Kievit et al., 2016; Kuznetsova et al., 2016; Penke et al., 2012; Schilling et al., 2023; Sexton et al., 2014; Voineskos et al., 2012; Weiss et al., 2024). It is worth noting though that FA and MSD also loaded appreciably on other WMFs, suggesting that these individual metrics are likely to capture multiple different aspects of WM health.

Interestingly, the fourth WMF related to microstructural complexity / WM volume and which made a unique contribution to fluid intelligence, loaded mainly on mean signal kurtosis (MSK). Kurtosis imaging captures the non-Gaussian diffusivity in WM tracts, and lower MSK values are thought to reflect reduced tissue complexity, as they indicate water diffusion is more Gaussian or unrestricted by the presence of neuron axons, dendrites and cells. It potentially provides a broader measure of microstructural integrity than DTI measures or the information captured by NDI. This is consistent with the idea that age-related processes lead to changes in multiple cell types in WM tracts, and these changes are best captured by a broad measure like MSK (Graciani et al., 2023). It may therefore be a useful measure for future studies of neurocognitive ageing.

Beside our main focus on the latent WMFs, we also examined the relationship between each individual WM metric and cognition (see supplementary materials). T1/T2, WMV and WMHI all showed strong age-related changes, but these measures did not show robust relationship with cognition after accounting for age effects. While the T1/T2 ratio has gained some popularity recently, these results suggest that it may not be a particularly useful measure to understand cognition. This is in line with recent work suggesting that the T1/T2 ratio in WM tracts is not particularly sensitive to myelin content and may be difficult to interpret (Arshad et al., 2017; Glasser et al., 2022; Sandrone et al., 2023; Uddin et al., 2018, 2019). On the other hand, given the large number of neuroimaging studies reporting an association between WMHI, age and cognition (Cox, Lyall, et al., 2019; d’Arbeloff et al., 2019; Debette & Markus, 2010; Freeze et al., 2020; Jacobs et al., 2013; Vergoossen et al., 2021), we were surprised to only see a weak association between WMHI and cognitive performance. However, there may be good reasons for this. First, we computed WMHI using T1w and T2w images, rather than the FLAIR images – which better capture the margins of white matter lesions. Second, we focused on a whole adult life-span sample, whereas most prior work on WMHI and cognition has focused on older adults, particularly those with cardiovascular disorders. Third, it may be important to consider the number and/or location of the WMHI lesions, rather than focusing on just their combined volume, as here (Biesbroek et al., 2016; Petersen et al., 2024).

There are other limitations to this work that are important to consider. The cross-sectional nature of the data prevented us from testing how longitudinal changes in WM relate to age-related cognitive changes (Lawrence et al., 2021; Raz & Lindenberger, 2011; Rodriguez-Ayllon et al., 2024; Walhovd et al., 2023). Secondly, we used linear latent variable methods such as PCA, but it is possible that there are non-linear associations between the different WM measures (e.g., see Farooq et al., 2024; Y. Raykov & Saad, 2022). Thirdly, we did not consider regional variation in WM across the brain, for reasons explained above. Lastly, in order to use comparable brain regions across metrics, we used the same set of atlas-based major WM tracts, inverse-normalised into the native space of each participant; it might be possible to extract more sensitive metrics by aligning participants to a WM skeleton (based on DTI) defined in each individual’s native space (as done, for example, for standard PSMD, Baykara et al., 2016).

In conclusion, we show that utilising multiple white matter measures from multiple MR contrasts, together with dimensionality reduction techniques, can lead to robust and interpretable white matter health measures. We show that 4 factors are sufficient to capture the majority of variance in those measures, and that these factors uniquely relate to cardiovascular and cognitive measures. We argue that this multidimensional approach to white matter health may be a fruitful avenue to better understand the links between aging, brain and cognition.

## Data and Code Availability

The raw data are in BIDS format are available on request from this website: https://camcan-archive.mrc-cbu.cam.ac.uk/dataaccess/. The pre-processed ROI and cognitive data and code to reproduce the main results from the pre-registered analyses are available here - https://osf.io/y7ct8/.

## Author Contributions

Petar P. Raykov: Methodology, Software, Validation, Formal analysis, Investigation, Data Curation, Writing—Original Draft, Writing—Review & Editing, Visualisation. Marta Morgado Correia: Conceptualisation, Methodology, Validation; Kamen Tsvetanov: Methodology, Validation; Rafael N. Henriques: Conceptualisation, Methodology, Validation, Data Curation; Alberto Del Cerro León: Methodology, Validation, Data Curation; Matthew Bracher-Smith: Methodology, Validation, Data Curation; Valentina Escott-Price: Methodology, Validation, Data Curation, Review & Editing; Yordan P. Raykov: Methodology, Validation; Richard N. Henson: Conceptualisation, Methodology, Supervision, Validation, Writing – Review & Editing, Resources, Project administration and Funding acquisition.

## Author Rights Retention

For the purpose of open access, the author has applied a Creative Commons Attribution (CC BY) licence to any Author Accepted Manuscript version arising from this submission.

## Ethics Statement

Approval for the Cam-CAN study was granted by the Research Ethics Committee of Cambridgeshire 2 (now known as East of England—Cambridge Central). Prior to their involvement, participants provided written, informed consent.

## Acknowledgements

Cam-CAN Project principal personnel: Richard N Henson, Lorraine K Tyler, Kamen A Tsvetanov, Carol Brayne, Edward T Bullmore, Andrew C Calder, Rhodri Cusack, Tim Dalgleish, John Duncan, Fiona E Matthews, William D Marslen-Wilson, James B Rowe, Meredith A Shafto; Research Associates: Karen Campbell, Teresa Cheung, Simon Davis, Linda Geerligs, Rogier Kievit, Anna McCarrey, Abdur Mustafa, Darren Price, David Samu, Jason R Taylor, Matthias Treder, Janna van Belle, Nitin Williams, Daniel Mitchell, Simon Fisher, Else Eising, Ethan Knights, Adam Attaheri; Research Assistants: Lauren Bates, Tina Emery, Sharon Erzinçlioğlu, Andrew Gadie, Sofia Gerbase, Stanimira Georgieva, Claire Hanley, Beth Parkin, David Troy, Ina Demetriou, Will Ducktt; Affiliated Personnel: Tibor Auer, Marta Correia, Lu Gao, Emma Green, Rafael Henriques; Research Interviewers: Jodie Allen, Gillian Amery, Liana Amunts, Anne Barcroft, Amanda Castle, Cheryl Dias, Jonathan Dowrick, Melissa Fair, Hayley Fisher, Anna Goulding, Adarsh Grewal, Geoff Hale, Andrew Hilton, Frances Johnson, Patricia Johnston, Thea Kavanagh-Williamson, Magdalena Kwasniewska, Alison McMinn, Kim Norman, Jessica Penrose, Fiona Roby, Diane Rowland, John Sargeant, Maggie Squire, Beth Stevens, Aldabra Stoddart, Cheryl Stone, Tracy Thompson, Ozlem Yazlik; and administrative staff: Dan Barnes, Marie Dixon, Jaya Hillman, Joanne Mitchell, Laura Villis.

## References

Arshad, M., Stanley, J. A., & Raz, N. (2017). Test–retest reliability and concurrent validity of in vivo myelin content indices: Myelin water fraction and calibrated T1w/T2w image ratio. Human Brain Mapping, 38(4), 1780–1790.

Ashburner, J. (2007). A fast diffeomorphic image registration algorithm. Neuroimage, 38(1), 95–113.

Basser, P. J., Mattiello, J., & LeBihan, D. (1994). MR diffusion tensor spectroscopy and imaging. Biophysical Journal, 66(1), 259–267.

Baykara, E., Gesierich, B., Adam, R., Tuladhar, A. M., Biesbroek, J. M., Koek, H. L., Ropele, S., Jouvent, E., Chabriat, H., Ertl-Wagner, B., Ewers, M., Schmidt, R., de Leeuw, F. E., Biessels, G. J., Dichgans, M., & Duering, M. (2016). A Novel Imaging Marker for Small Vessel Disease Based on Skeletonization of White Matter Tracts and Diffusion Histograms. Annals of Neurology. 10.1002/ana.24758

Biesbroek, J. M., Weaver, N. A., Hilal, S., Kuijf, H. J., Ikram, M. K., Xu, X., Tan, B. Y., Venketasubramanian, N., Postma, A., & Biessels, G. J. (2016). Impact of strategically located white matter hyperintensities on cognition in memory clinic patients with small vessel disease. PLoS One, 11(11), e0166261.

Black, S., Gao, F., & Bilbao, J. (2009). Understanding white matter disease: Imaging-pathological correlations in vascular cognitive impairment. Stroke, 40(3_suppl_1), S48–S52.

Boaventura, M., Sastre-Garriga, J., Rimkus, C. de M., Rovira, À., & Pareto, D. (2024). T1/T2-weighted ratio: A feasible MRI biomarker in multiple sclerosis. Multiple Sclerosis Journal, 13524585241233448.

Boroshok, A. L., McDermott, C. L., Fotiadis, P., Park, A. T., Tooley, U. A., Gataviņš, M. M., Tisdall, M. D., Bassett, D. S., & Mackey, A. P. (2023). Individual differences in T1w/T2w ratio development during childhood. Developmental Cognitive Neuroscience, 62, 101270.

Botz, J., Lohner, V., & Schirmer, M. D. (2023). Spatial patterns of white matter hyperintensities: A systematic review. Frontiers in Aging Neuroscience, 15, 1165324.

Bro, R., Kjeldahl, K., Smilde, A. K., & Kiers, H. A. L. (2008). Cross-validation of component models: A critical look at current methods. Analytical and Bioanalytical Chemistry, 390, 1241–1251.

Camacho, J., & Ferrer, A. (2012). Cross-validation in PCA models with the element-wise k-fold (ekf) algorithm: Theoretical aspects. Journal of Chemometrics, 26(7), 361–373.

Camacho, J., Pérez-Villegas, A., Rodríguez-Gómez, R. A., & Jiménez-Mañas, E. (2015). Multivariate exploratory data analysis (MEDA) toolbox for Matlab. Chemometrics and Intelligent Laboratory Systems, 143, 49–57.

Cattell, R. B. (1971). Abilities: Their structure, growth, and action. In Abilities: Their structure, growth, and action. (pp. xxii, 583–xxii, 583). Houghton Mifflin.

Cerri, S., Greve, D. N., Hoopes, A., Lundell, H., Siebner, H. R., Mühlau, M., & Van Leemput, K. (2023). An open-source tool for longitudinal whole-brain and white matter lesion segmentation. NeuroImage: Clinical, 38, 103354.

Chamberland, M., Raven, E. P., Genc, S., Duffy, K., Descoteaux, M., Parker, G. D., Tax, C. M., & Jones, D. K. (2019). Dimensionality reduction of diffusion MRI measures for improved tractometry of the human brain. NeuroImage, 200, 89–100.

Corley, J., Conte, F., Harris, S. E., Taylor, A. M., Redmond, P., Russ, T. C., Deary, I. J., & Cox, S. R. (2023). Predictors of longitudinal cognitive ageing from age 70 to 82 including APOE e4 status, early-life and lifestyle factors: The Lothian Birth Cohort 1936. Molecular Psychiatry, 28(3), 1256–1271.

Cox, S. R., Lyall, D. M., Ritchie, S. J., Bastin, M. E., Harris, M. A., Buchanan, C. R., Fawns-Ritchie, C., Barbu, M. C., De Nooij, L., & Reus, L. M. (2019). Associations between vascular risk factors and brain MRI indices in UK Biobank. European Heart Journal, 40(28), 2290–2300.

Cox, S. R., Ritchie, S. J., Fawns-Ritchie, C., Tucker-Drob, E. M., & Deary, I. J. (2019). Structural brain imaging correlates of general intelligence in UK Biobank. Intelligence, 76, 101376.

Cox, S. R., Ritchie, S. J., Tucker-Drob, E. M., Liewald, D. C., Hagenaars, S. P., Davies, G., Wardlaw, J. M., Gale, C. R., Bastin, M. E., & Deary, I. J. (2016). Ageing and brain white matter structure in 3,513 UK Biobank participants. Nature Communications, 7(1), 13629.

d’Arbeloff, T., Elliott, M. L., Knodt, A. R., Melzer, T. R., Keenan, R., Ireland, D., Ramrakha, S., Poulton, R., Anderson, T., & Caspi, A. (2019). White matter hyperintensities are common in midlife and already associated with cognitive decline. Brain Communications, 1(1), fcz041.

d’Arbeloff, T., Elliott, M. L., Knodt, A. R., Sison, M., Melzer, T. R., Ireland, D., Ramrakha, S., Poulton, R., Caspi, A., & Moffitt, T. E. (2021). Midlife cardiovascular fitness is reflected in the Brain’s White matter. Frontiers in Aging Neuroscience, 13, 652575.

Daskalopoulou, C., Stubbs, B., Kralj, C., Koukounari, A., Prince, M., & Prina, A. M. (2017). Physical activity and healthy ageing: A systematic review and meta-analysis of longitudinal cohort studies. Ageing Research Reviews, 38, 6–17.

Davis, S. W., Dennis, N. A., Buchler, N. G., White, L. E., Madden, D. J., & Cabeza, R. (2009). Assessing the effects of age on long white matter tracts using diffusion tensor tractography. Neuroimage, 46(2), 530–541.

Deary, I. J., Corley, J., Gow, A. J., Harris, S. E., Houlihan, L. M., Marioni, R. E., Penke, L., Rafnsson, S. B., & Starr, J. M. (2009). Age-associated cognitive decline. British Medical Bulletin, 92(1), 135–152.

Deary, I. J., Penke, L., & Johnson, W. (2010). The neuroscience of human intelligence differences. Nature Reviews Neuroscience, 11(3), 201–211.

Debette, S., & Markus, H. S. (2010). The clinical importance of white matter hyperintensities on brain magnetic resonance imaging: Systematic review and meta-analysis. Bmj, 341.

Dekkers, I. A., Jansen, P. R., & Lamb, H. J. (2019). Obesity, brain volume, and white matter microstructure at MRI: a cross-sectional UK biobank study. Radiology, 291(3), 763–771.

Demirtaş, M., Burt, J. B., Helmer, M., Ji, J. L., Adkinson, B. D., Glasser, M. F., Van Essen, D. C., Sotiropoulos, S. N., Anticevic, A., & Murray, J. D. (2019). Hierarchical heterogeneity across human cortex shapes large-scale neural dynamics. Neuron, 101(6), 1181–1194.

Farooq, A., Raykov, Y. P., Raykov, P., & Little, M. A. (2024). Adaptive Latent Feature Sharing for Piecewise Linear Dimensionality Reduction. Journal of Machine Learning Research, 25(135), 1–42.

Folstein, M. F., Folstein, S. E., & McHugh, P. R. (1975). “Mini-mental state”: A practical method for grading the cognitive state of patients for the clinician. Journal of Psychiatric Research, 12(3), 189–198.

Freeze, W. M., Jacobs, H. I. L., De Jong, J. J., Verheggen, I. C. M., Gronenschild, E. H. B. M., Palm, W. M., Hoff, E. I., Wardlaw, J. M., Jansen, J. F. A., & Verhey, F. R. (2020). White matter hyperintensities mediate the association between blood-brain barrier leakage and information processing speed. Neurobiology of Aging, 85, 113–122.

Fuhrmann, D., Nesbitt, D., Shafto, M., Rowe, J. B., Price, D., Gadie, A., Tyler, L. K., Brayne, C., Bullmore, E. T., & Calder, A. C. (2019). Strong and specific associations between cardiovascular risk factors and white matter micro-and macrostructure in healthy aging. Neurobiology of Aging, 74, 46–55.

Ganzetti, M., Wenderoth, N., & Mantini, D. (2014). Whole brain myelin mapping using T1-and T2-weighted MR imaging data. Frontiers in Human Neuroscience, 8, 671.

Ge, T., Chen, C.-Y., Ni, Y., Feng, Y.-C. A., & Smoller, J. W. (2019). Polygenic prediction via Bayesian regression and continuous shrinkage priors. Nature Communications, 10(1), 1776.

Glasser, M. F., Coalson, T. S., Harms, M. P., Xu, J., Baum, G. L., Autio, J. A., Auerbach, E. J., Greve, D. N., Yacoub, E., & Van Essen, D. C. (2022). Empirical transmit field bias correction of T1w/T2w myelin maps. NeuroImage, 258, 119360.

Glasser, M. F., & Van Essen, D. C. (2011). Mapping human cortical areas in vivo based on myelin content as revealed by T1-and T2-weighted MRI. Journal of Neuroscience, 31(32), 11597–11616.

Graciani, A. L., Gutierre, M. U., Coppi, A. A., Arida, R. M., & Gutierre, R. C. (2023). Myelin, aging, and physical exercise. Neurobiology of Aging, 127, 70–81.

Graham, J. W. (2003). Adding missing-data-relevant variables to FIML-based structural equation models. Structural Equation Modeling, 10(1), 80–100.

Grieve, S. M., Williams, L. M., Paul, R. H., Clark, C. R., & Gordon, E. (2007). Cognitive aging, executive function, and fractional anisotropy: A diffusion tensor MR imaging study. American Journal of Neuroradiology, 28(2), 226–235.

Grydeland, H., Vértes, P. E., Váša, F., Romero-Garcia, R., Whitaker, K., Alexander-Bloch, A. F., Bjørnerud, A., Patel, A. X., Sederevičius, D., & Tamnes, C. K. (2019). Waves of maturation and senescence in micro-structural MRI markers of human cortical myelination over the lifespan. Cerebral Cortex, 29(3), 1369–1381.

Grydeland, H., Walhovd, K. B., Tamnes, C. K., Westlye, L. T., & Fjell, A. M. (2013). Intracortical myelin links with performance variability across the human lifespan: Results from T1-and T2-weighted MRI myelin mapping and diffusion tensor imaging. Journal of Neuroscience, 33(47), 18618–18630.

Gunning-Dixon, F. M., Brickman, A. M., Cheng, J. C., & Alexopoulos, G. S. (2009). Aging of cerebral white matter: A review of MRI findings. International Journal of Geriatric Psychiatry: A Journal of the Psychiatry of Late Life and Allied Sciences, 24(2), 109–117.

Henriques, R. N. (2018). Advanced methods for diffusion MRI data analysis and their application to the healthy ageing brain.

Henriques, R. N., Correia, M. M., Marrale, M., Huber, E., Kruper, J., Koudoro, S., Yeatman, J. D., Garyfallidis, E., & Rokem, A. (2021a). Diffusional kurtosis imaging in the diffusion imaging in python project. Frontiers in Human Neuroscience, 15, 675433.

Henriques, R. N., Correia, M. M., Marrale, M., Huber, E., Kruper, J., Koudoro, S., Yeatman, J. D., Garyfallidis, E., & Rokem, A. (2021b). Diffusional kurtosis imaging in the diffusion imaging in python project. Frontiers in Human Neuroscience, 15, 675433.

Henriques, R. N., Henson, R., & Correia, M. M. (2023). Unique information from common diffusion MRI models about white-matter differences across the human adult lifespan. ArXiv Preprint ArXiv:2306.09942.

Henriques, R. N., Jespersen, S. N., Jones, D. K., & Veraart, J. (2021). Toward more robust and reproducible diffusion kurtosis imaging. Magnetic Resonance in Medicine, 86(3), 1600–1613.

Henson, R. N., Suri, S., Knights, E., Rowe, J. B., Kievit, R. A., Lyall, D. M., Chan, D., Eising, E., & Fisher, S. E. (2020). Effect of apolipoprotein E polymorphism on cognition and brain in the Cambridge Centre for Ageing and Neuroscience cohort. Brain and Neuroscience Advances, 4, 2398212820961704.

Hernández, M. del C. V., Booth, T., Murray, C., Gow, A. J., Penke, L., Morris, Z., Maniega, S. M., Royle, N. A., Aribisala, B. S., & Bastin, M. E. (2013). Brain white matter damage in aging and cognitive ability in youth and older age. Neurobiology of Aging, 34(12), 2740–2747.

Hofman, A., Rodriguez-Ayllon, M., Vernooij, M. W., Croll, P. H., Luik, A. I., Neumann, A., Niessen, W. J., Ikram, M. A., Voortman, T., & Muetzel, R. L. (2023). Physical activity levels and brain structure in middle-aged and older adults: A bidirectional longitudinal population-based study. Neurobiology of Aging, 121, 28–37.

Huber, E., Henriques, R. N., Owen, J. P., Rokem, A., & Yeatman, J. D. (2019). Applying microstructural models to understand the role of white matter in cognitive development. Developmental Cognitive Neuroscience, 36, 100624.

Jacobs, H. I. L., Leritz, E. C., Williams, V. J., Van Boxtel, M. P. J., Elst, W. van der, Jolles, J., Verhey, F. R. J., McGlinchey, R. E., Milberg, W. P., & Salat, D. H. (2013). Association between white matter microstructure, executive functions, and processing speed in older adults: The impact of vascular health. Human Brain Mapping, 34(1), 77–95.

Jenkinson, M., Beckmann, C. F., Behrens, T. E. J., Woolrich, M. W., & Smith, S. M. (2012). Fsl. Neuroimage, 62(2), 782–790.

Jensen, J. H., & Helpern, J. A. (2010). MRI quantification of non-Gaussian water diffusion by kurtosis analysis. NMR in Biomedicine, 23(7), 698–710.

Jensen, J. H., Helpern, J. A., Ramani, A., Lu, H., & Kaczynski, K. (2005). Diffusional kurtosis imaging: The quantification of non-gaussian water diffusion by means of magnetic resonance imaging. Magnetic Resonance in Medicine: An Official Journal of the International Society for Magnetic Resonance in Medicine, 53(6), 1432–1440.

Josse, J., & Husson, F. (2012). Selecting the number of components in principal component analysis using cross-validation approximations. Computational Statistics & Data Analysis, 56(6), 1869–1879.

Kamiya, K., Kamagata, K., Ogaki, K., Hatano, T., Ogawa, T., Takeshige-Amano, H., Murata, S., Andica, C., Murata, K., & Feiweier, T. (2020). Brain white-matter degeneration due to aging and Parkinson disease as revealed by double diffusion encoding. Frontiers in Neuroscience, 14, 584510.

Kang, P., Ying, C., Chen, Y., Ford, A. L., An, H., & Lee, J.-M. (2022). Oxygen metabolic stress and white matter injury in patients with cerebral small vessel disease. Stroke, 53(5), 1570–1579.

Kellner, E., Dhital, B., Kiselev, V. G., & Reisert, M. (2016). Gibbs-ringing artifact removal based on local subvoxel-shifts. Magnetic Resonance in Medicine, 76(5), 1574–1581.

Kennedy, K. M., & Raz, N. (2009). Pattern of normal age-related regional differences in white matter microstructure is modified by vascular risk. Brain Research, 1297, 41–56.

Khormi, I., Al-Iedani, O., Alshehri, A., Ramadan, S., & Lechner-Scott, J. (2023). MR myelin imaging in multiple sclerosis: A scoping review. Journal of the Neurological Sciences, 122807.

Kievit, R. A., Davis, S. W., Griffiths, J., Correia, M. M., & Henson, R. N. (2016). A watershed model of individual differences in fluid intelligence. Neuropsychologia, 91, 186–198.

King, D. L. O., Henson, R. N., Correia, M. M., Rowe, J. B., & Tsvetanov, K. A. (2024). PULSE PRESSURE IMPAIRS COGNITION VIA WHITE MATTER DISRUPTION. MedRxiv, 2024–12.

King, D. L. O., Henson, R. N., Kievit, R., Wolpe, N., Brayne, C., Tyler, L. K., Rowe, J. B., & Tsvetanov, K. A. (2023). Distinct components of cardiovascular health are linked with age-related differences in cognitive abilities. Scientific Reports, 13(1), 978.

Kuznetsova, K. A., Maniega, S. M., Ritchie, S. J., Cox, S. R., Storkey, A. J., Starr, J. M., Wardlaw, J. M., Deary, I. J., & Bastin, M. E. (2016). Brain white matter structure and information processing speed in healthy older age. Brain Structure and Function, 221, 3223–3235.

Lakatta, E. G., & Levy, D. (2003). Arterial and cardiac aging: Major shareholders in cardiovascular disease enterprises: Part I: aging arteries: A “set up” for vascular disease. Circulation, 107(1), 139–146.

Lawrence, K. E., Nabulsi, L., Santhalingam, V., Abaryan, Z., Villalon-Reina, J. E., Nir, T. M., Ba Gari, I., Zhu, A. H., Haddad, E., & Muir, A. M. (2021). Age and sex effects on advanced white matter microstructure measures in 15,628 older adults: A UK biobank study. Brain Imaging and Behavior, 15(6), 2813–2823.

Le Bihan, D., & Johansen-Berg, H. (2012). Diffusion MRI at 25: Exploring brain tissue structure and function. Neuroimage, 61(2), 324–341.

Lebel, C., Gee, M., Camicioli, R., Wieler, M., Martin, W., & Beaulieu, C. (2012). Diffusion tensor imaging of white matter tract evolution over the lifespan. Neuroimage, 60(1), 340–352.

Lee, S.-N., Woo, S.-H., Lee, E. J., Kim, K. K., & Kim, H.-R. (2024). Association between T1w/T2w ratio in white matter and cognitive function in Alzheimer’s disease. Scientific Reports, 14(1), 7228.

Little, R. J., & Rubin, D. B. (2019). Statistical analysis with missing data (Vol. 793). John Wiley & Sons.

Liu, X., Tyler, L. K., Davis, S. W., Rowe, J. B., & Tsvetanov, K. A. (2023). Cognition’s dependence on functional network integrity with age is conditional on structural network integrity. Neurobiology of Aging.

Livingston, G., Huntley, J., Sommerlad, A., Ames, D., Ballard, C., Banerjee, S., Brayne, C., Burns, A., Cohen-Mansfield, J., Cooper, C., Costafreda, S. G., Dias, A., Fox, N., Gitlin, L. N., Howard, R., Kales, H. C., Kivimäki, M., Larson, E. B., Ogunniyi, A., … Mukadam, N. (2020). Dementia prevention, intervention, and care: 2020 report of the *Lancet* Commission. The Lancet, 396(10248), 413–446. 10.1016/S0140-6736(20)30367-6

Lyall, D. M., Cox, S. R., Lyall, L. M., Celis-Morales, C., Cullen, B., Mackay, D. F., Ward, J., Strawbridge, R. J., McIntosh, A. M., & Sattar, N. (2020). Association between APOE e4 and white matter hyperintensity volume, but not total brain volume or white matter integrity. Brain Imaging and Behavior, 14, 1468–1476.

Maillard, P., Carmichael, O., Harvey, D., Fletcher, E., Reed, B., Mungas, D., & DeCarli, C. (2013). FLAIR and diffusion MRI signals are independent predictors of white matter hyperintensities. American Journal of Neuroradiology, 34(1), 54–61.

Mancini, M., Karakuzu, A., Cohen-Adad, J., Cercignani, M., Nichols, T. E., & Stikov, N. (2020). An interactive meta-analysis of MRI biomarkers of myelin. Elife, 9, e61523.

Martinez Sosa, S., & Smith, K. J. (2017). Understanding a role for hypoxia in lesion formation and location in the deep and periventricular white matter in small vessel disease and multiple sclerosis. Clinical Science, 131(20), 2503–2524.

Mendez Colmenares, A., Prytherch, B., Thomas, M. L., & Burzynska, A. Z. (2023). Within-person changes in the aging white matter microstructure and their modifiers: A meta-analysis and systematic review of longitudinal diffusion tensor imaging studies. Imaging Neuroscience, 1, 1–32.

Moon, S. Y., de Souto Barreto, P., Cesari, M., Chupin, M., Mangin, J. F., Bouyahia, A., Fillon, L., Andrieu, S., & Vellas, B. (2018). Physical activity and changes in white matter hyperintensities over three years. *The Journal of Nutrition*, Health and Aging, 22(3), 425–430.

Moreno-Agostino, D., Daskalopoulou, C., Wu, Y.-T., Koukounari, A., Haro, J. M., Tyrovolas, S., Panagiotakos, D. B., Prince, M., & Prina, A. M. (2020). The impact of physical activity on healthy ageing trajectories: Evidence from eight cohort studies. International Journal of Behavioral Nutrition and Physical Activity, 17, 1–12.

Moseley, M. (2002). Diffusion tensor imaging and aging–a review. NMR in Biomedicine: An International Journal Devoted to the Development and Application of Magnetic Resonance In Vivo, 15(7-8), 553–560.

Novikov, D. S., Kiselev, V. G., & Jespersen, S. N. (2018). On modeling. Magnetic Resonance in Medicine, 79(6), 3172–3193.

Nusbaum, A. O., Tang, C. Y., Buchsbaum, M. S., Wei, T. C., & Atlas, S. W. (2001). Regional and global changes in cerebral diffusion with normal aging. American Journal of Neuroradiology, 22(1), 136–142.

Odrobina, E. E., Lam, T. Y. J., Pun, T., Midha, R., & Stanisz, G. J. (2005). MR properties of excised neural tissue following experimentally induced demyelination. NMR in Biomedicine, 18(5), 277–284.

Palta, P., Sharrett, A. R., Gabriel, K. P., Gottesman, R. F., Folsom, A. R., Power, M. C., Evenson, K. R., Jack Jr, C. R., Knopman, D. S., & Mosley, T. H. (2021). Prospective analysis of leisure-time physical activity in midlife and beyond and brain damage on MRI in older adults. Neurology, 96(7), e964–e974.

Pannese, E. (2011). Morphological changes in nerve cells during normal aging. Brain Structure and Function, 216(2), 85–89.

Pantoni, L., Garcia, J. H., & Gutierrez, J. A. (1996). Cerebral white matter is highly vulnerable to ischemia. Stroke, 27(9), 1641–1647.

Pelkmans, W., Dicks, E., Barkhof, F., Vrenken, H., Scheltens, P., van der Flier, W. M., & Tijms, B. M. (2019). Gray matter T1-w/T2-w ratios are higher in Alzheimer’s disease. Human Brain Mapping, 40(13), 3900–3909.

Pelletier, A., Bernard, C., Dilharreguy, B., Helmer, C., Le Goff, M., Chanraud, S., Dartigues, J.-F., Allard, M., Amieva, H., & Catheline, G. (2017). Patterns of brain atrophy associated with episodic memory and semantic fluency decline in aging. Aging (Albany NY), 9(3), 741.

Penke, L., Maniega, S. M., Bastin, M. E., Hernandez, V., Murray, C., Royle, N. A., Starr, J. M., Wardlaw, J. M., & Deary, I. J. (2012). Brain white matter tract integrity as a neural foundation for general intelligence. Molecular Psychiatry, 17(10), 1026–1030.

Petersen, M., Coenen, M., DeCarli, C., De Luca, A., van der Lelij, E., Barkhof, F., Benke, T., Chen, C. P., Dal-Bianco, P., & Dewenter, A. (2024). Enhancing cognitive performance prediction by white matter hyperintensity connectivity assessment. *Brain*, awae315.

Prins, N. D., & Scheltens, P. (2015). White matter hyperintensities, cognitive impairment and dementia: An update. Nature Reviews Neurology, 11(3), 157–165.

Raichlen, D. A., Ally, M., Aslan, D. H., Sayre, M. K., Bharadwaj, P. K., Maltagliati, S., Lai, M. H. C., Wilcox, R. R., Habeck, C. G., & Klimentidis, Y. C. (2024). Associations between accelerometer-derived sedentary behavior and physical activity with white matter hyperintensities in middle-aged to older adults. Alzheimer’s & Dementia: Diagnosis, Assessment & Disease Monitoring, 16(3), e70001.

Ratcliff, R. (1993). Methods for dealing with reaction time outliers. Psychological Bulletin, 114(3), 510.

Raykov, P. P., Daly, J., Fisher, S. E., Eising, E., Geerligs, L., & Bird, C. M. (2025). No effect of apolipoprotein E polymorphism on MRI brain activity during movie watching. Brain and Neuroscience Advances, 9, 23982128251314576.

Raykov, Y., & Saad, D. (2022). Principled machine learning. IEEE Journal of Selected Topics in Quantum Electronics, 28(4: Mach. Learn. in Photon. Commun. and Meas. Syst.), 1–19.

Raz, N., & Lindenberger, U. (2011). Only time will tell: Cross-sectional studies offer no solution to the age–brain–cognition triangle: Comment on Salthouse (2011).

Reas, E. T., Laughlin, G. A., Hagler Jr, D. J., Lee, R. R., Dale, A. M., & McEvoy, L. K. (2021). Age and sex differences in the associations of pulse pressure with white matter and subcortical microstructure. Hypertension, 77(3), 938–947.

Reese, T. G., Heid, O., Weisskoff, R. M., & Wedeen, V. J. (2003). Reduction of eddy-current-induced distortion in diffusion MRI using a twice-refocused spin echo. Magnetic Resonance in Medicine: An Official Journal of the International Society for Magnetic Resonance in Medicine, 49(1), 177–182.

Ritchie, C. W., Bridgeman, K., Gregory, S., O’Brien, J. T., Danso, S. O., Dounavi, M.-E., Carriere, I., Driscoll, D., Hillary, R., & Koychev, I. (2024). The PREVENT Dementia programme: Baseline demographic, lifestyle, imaging and cognitive data from a midlife cohort study investigating risk factors for dementia. Brain Communications, 6(3).

Rodriguez-Ayllon, M., Neumann, A., Hofman, A., Vernooij, M. W., & Neitzel, J. (2024). The bidirectional relationship between brain structure and physical activity: A longitudinal analysis in the UK Biobank. Neurobiology of Aging, 138, 1–9.

Rosseel, Y. (2012). lavaan: An R package for structural equation modeling. Journal of Statistical Software, 48, 1–36.

Rowbotham, G., & Little, E. (1965). Circulations of the cerebral hemispheres. British Journal of Surgery, 52(1), 8–21.

Saccenti, E., & Camacho, J. (2015a). Determining the number of components in principal components analysis: A comparison of statistical, crossvalidation and approximated methods. Chemometrics and Intelligent Laboratory Systems, 149, 99–116.

Saccenti, E., & Camacho, J. (2015b). On the use of the observation-wise k-fold operation in PCA cross-validation. Journal of Chemometrics, 29(8), 467–478.

Sandrone, S., Aiello, M., Cavaliere, C., Thiebaut de Schotten, M., Reimann, K., Troakes, C., Bodi, I., Lacerda, L., Monti, S., & Murphy, D. (2023). Mapping myelin in white matter with T1-weighted/T2-weighted maps: Discrepancy with histology and other myelin MRI measures. Brain Structure and Function, 228(2), 525–535.

Savage, J. E., Jansen, P. R., Stringer, S., Watanabe, K., Bryois, J., De Leeuw, C. A., Nagel, M., Awasthi, S., Barr, P. B., & Coleman, J. R. (2018). Genome-wide association meta-analysis in 269,867 individuals identifies new genetic and functional links to intelligence. Nature Genetics, 50(7), 912–919.

Schilling, K. G., Archer, D., Yeh, F.-C., Rheault, F., Cai, L. Y., Shafer, A., Resnick, S. M., Hohman, T., Jefferson, A., & Anderson, A. W. (2023). Short superficial white matter and aging: A longitudinal multi-site study of 1293 subjects and 2711 sessions. Aging Brain, 3, 100067.

Sexton, C. E., Walhovd, K. B., Storsve, A. B., Tamnes, C. K., Westlye, L. T., Johansen-Berg, H., & Fjell, A. M. (2014). Accelerated changes in white matter microstructure during aging: A longitudinal diffusion tensor imaging study. Journal of Neuroscience, 34(46), 15425–15436.

Shafee, R., Buckner, R. L., & Fischl, B. (2015). Gray matter myelination of 1555 human brains using partial volume corrected MRI images. Neuroimage, 105, 473–485.

Shafto, M. A., Tyler, L. K., Dixon, M., Taylor, J. R., Rowe, J. B., Cusack, R., Calder, A. J., Marslen-Wilson, W. D., Duncan, J., & Dalgleish, T. (2014). The Cambridge Centre for Ageing and Neuroscience (Cam-CAN) study protocol: A cross-sectional, lifespan, multidisciplinary examination of healthy cognitive ageing. BMC Neurology, 14, 1–25.

Sled, J. G. (2018). Modelling and interpretation of magnetization transfer imaging in the brain. Neuroimage, 182, 128–135.

Smith, S. M., Jenkinson, M., Woolrich, M. W., Beckmann, C. F., Behrens, T. E. J., Johansen-Berg, H., Bannister, P. R., De Luca, M., Drobnjak, I., & Flitney, D. E. (2004). Advances in functional and structural MR image analysis and implementation as FSL. NeuroImage, 23, S208–S219. 10.1016/j.neuroimage.2004.07.051

Sotiropoulos, S. N., & Zalesky, A. (2019). Building connectomes using diffusion MRI: why, how and but. NMR in Biomedicine, 32(4), e3752.

Stanisz, G. J., Webb, S., Munro, C. A., Pun, T., & Midha, R. (2004). MR properties of excised neural tissue following experimentally induced inflammation. Magnetic Resonance in Medicine: An Official Journal of the International Society for Magnetic Resonance in Medicine, 51(3), 473–479.

Strömmer, J. M., Davis, S. W., Henson, R. N., Tyler, L. K., & Campbell, K. L. (2020). Physical activity predicts population-level age-related differences in frontal white matter. The Journals of Gerontology: Series A, 75(2), 236–243.

Sui, Y. V., Masurkar, A. V., Rusinek, H., Reisberg, B., & Lazar, M. (2022). Cortical myelin profile variations in healthy aging brain: A T1w/T2w ratio study. NeuroImage, 264, 119743.

Taylor, J. R., Williams, N., Cusack, R., Auer, T., Shafto, M. A., Dixon, M., Tyler, L. K., Cam-CAN, & Henson, R. N. (2017). The Cambridge Centre for Ageing and Neuroscience (Cam-CAN) data repository: Structural and functional MRI, MEG, and cognitive data from a cross-sectional adult lifespan sample. NeuroImage, 144, 262–269. 10.1016/j.neuroimage.2015.09.018

Thériault, R. (2023). rempsyc: Convenience functions for psychology. Journal of Open Source Software, 8(87), 5466.

Torres, E. R., Strack, E. F., Fernandez, C. E., Tumey, T. A., & Hitchcock, M. E. (2015). Physical activity and white matter hyperintensities: A systematic review of quantitative studies. Preventive Medicine Reports, 2, 319–325.

Torres-Simon, L., Cuesta, P., del Cerro-Leon, A., Chino, B., Orozco, L. H., Marsh, E. B., Gil, P., & Maestu, F. (2023). The effects of white matter hyperintensities on MEG power spectra in population with mild cognitive impairment. Frontiers in Human Neuroscience, 17, 1068216.

Torres-Simon, L., del Cerro-León, A., Yus, M., Bruña, R., Gil-Martinez, L., Marcos Dolado, A., Maestú, F., Arrazola-Garcia, J., & Cuesta, P. (2024). Decoding the Best Automated Segmentation Tools for Vascular White Matter Hyperintensities in the Aging Brain: A Clinician’s Guide to Precision and Purpose. MedRxiv, 2023.03.30.23287946. 10.1101/2023.03.30.23287946

Tsvetanov, K. A., Henson, R. N. A., Tyler, L. K., Razi, A., Geerligs, L., Ham, T. E., & Rowe, J. B. (2016). Extrinsic and intrinsic brain network connectivity maintains cognition across the lifespan despite accelerated decay of regional brain activation. Journal of Neuroscience, 36(11), 3115–3126.

Uddin, M. N., Figley, T. D., Marrie, R. A., Figley, C. R., & Group, C. S. (2018). Can T1w/T2w ratio be used as a myelin-specific measure in subcortical structures? Comparisons between FSE-based T1w/T2w ratios, GRASE-based T1w/T2w ratios and multi-echo GRASE-based myelin water fractions. NMR in Biomedicine, 31(3), e3868.

Uddin, M. N., Figley, T. D., Solar, K. G., Shatil, A. S., & Figley, C. R. (2019). Comparisons between multi-component myelin water fraction, T1w/T2w ratio, and diffusion tensor imaging measures in healthy human brain structures. Scientific Reports, 9(1), 2500.

Vavasour, I. M., Laule, C., Li, D. K. B., Traboulsee, A. L., & MacKay, A. L. (2011). Is the magnetization transfer ratio a marker for myelin in multiple sclerosis? Journal of Magnetic Resonance Imaging, 33(3), 710–718.

Veraart, J., Novikov, D. S., Christiaens, D., Ades-Aron, B., Sijbers, J., & Fieremans, E. (2016). Denoising of diffusion MRI using random matrix theory. Neuroimage, 142, 394–406.

Vergoossen, L. W. M., Jansen, J. F. A., van Sloten, T. T., Stehouwer, C. D. A., Schaper, N. C., Wesselius, A., Dagnelie, P. C., Köhler, S., van Boxtel, M. P. J., & Kroon, A. A. (2021). Interplay of white matter hyperintensities, cerebral networks, and cognitive function in an adult population: Diffusion-tensor imaging in the maastricht study. Radiology, 298(2), 384–392.

Vidal-Piñeiro, D., Walhovd, K. B., Storsve, A. B., Grydeland, H., Rohani, D. A., & Fjell, A. M. (2016). Accelerated longitudinal gray/white matter contrast decline in aging in lightly myelinated cortical regions. Human Brain Mapping, 37(10), 3669–3684.

Voineskos, A. N., Rajji, T. K., Lobaugh, N. J., Miranda, D., Shenton, M. E., Kennedy, J. L., Pollock, B. G., & Mulsant, B. H. (2012). Age-related decline in white matter tract integrity and cognitive performance: A DTI tractography and structural equation modeling study. Neurobiology of Aging, 33(1), 21–34.

Walhovd, K., Lövden, M., & Fjell, A. (2023). Timing of lifespan influences on brain and cognition. *Trends in Cognitive Sciences*, S1364–6613.

Wartolowska, K. A., & Webb, A. J. (2022). White matter damage due to pulsatile versus steady blood pressure differs by vascular territory: A cross-sectional analysis of the UK Biobank cohort study. Journal of Cerebral Blood Flow & Metabolism, 42(5), 802–810.

Wartolowska, K. A., & Webb, A. J. S. (2021). Midlife blood pressure is associated with the severity of white matter hyperintensities: Analysis of the UK Biobank cohort study. European Heart Journal, 42(7), 750–757.

Wei, J., Palta, P., Meyer, M. L., Kucharska-Newton, A., Pence, B. W., Aiello, A. E., Power, M. C., Walker, K. A., Sharrett, A. R., & Tanaka, H. (2020). Aortic stiffness and white matter microstructural integrity assessed by diffusion tensor imaging: The ARIC-NCS. Journal of the American Heart Association, 9(6), e014868.

Weiss, J., Beydoun, M. A., Beydoun, H. A., Georgescu, M. F., Hu, Y.-H., Hooten, N. N., Banerjee, S., Launer, L. J., Evans, M. K., & Zonderman, A. B. (2024). Pathways explaining racial/ethnic and socio-economic disparities in brain white matter integrity outcomes in the UK Biobank study. SSM-Population Health, 26, 101655.

Westlye, L. T., Walhovd, K. B., Dale, A. M., Bjørnerud, A., Due-Tønnessen, P., Engvig, A., Grydeland, H., Tamnes, C. K., Østby, Y., & Fjell, A. M. (2010). Life-span changes of the human brain white matter: Diffusion tensor imaging (DTI) and volumetry. Cerebral Cortex, 20(9), 2055–2068.

White, N. S., Leergaard, T. B., D’Arceuil, H., Bjaalie, J. G., & Dale, A. M. (2013). Probing tissue microstructure with restriction spectrum imaging: Histological and theoretical validation. Human Brain Mapping, 34(2), 327–346.

Winzeck, S. (2021). Methods for Data Management in Multi-Centre MRI Studies and Applications to Traumatic Brain Injury.

Woolrich, M. W., Jbabdi, S., Patenaude, B., Chappell, M., Makni, S., Behrens, T., Beckmann, C., Jenkinson, M., & Smith, S. M. (2009). Bayesian analysis of neuroimaging data in FSL. Neuroimage, 45(1), S173–S186.

Wozniak, J. R., & Lim, K. O. (2006). Advances in white matter imaging: A review of in vivo magnetic resonance methodologies and their applicability to the study of development and aging. Neuroscience & Biobehavioral Reviews, 30(6), 762–774.

Yeatman, J. D., Wandell, B. A., & Mezer, A. A. (2014). Lifespan maturation and degeneration of human brain white matter. Nature Communications, 5(1), 4932.

Zhang, H., Schneider, T., Wheeler-Kingshott, C. A., & Alexander, D. C. (2012). NODDI: practical in vivo neurite orientation dispersion and density imaging of the human brain. Neuroimage, 61(4), 1000–1016.

Zheng, Y., Lee, J., Rudick, R., & Fisher, E. (2018). Long-term magnetization transfer ratio evolution in multiple sclerosis white matter lesions. Journal of Neuroimaging, 28(2), 191–198.

