## Supplementary file for "Complementary MR measures of white matter and their relation to cardiovascular health and cognition"

### Supplementary Materials

#### Factor Analysis of cardiovascular measures

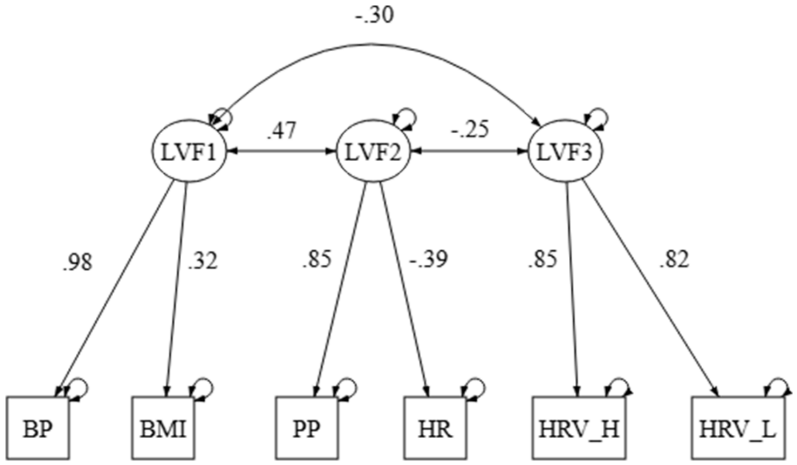


**Supplementary Figure 1** *Latent Cardiovascular measures.* Figure shows main loadings for the three cardio vascular latent factors. Here we followed a previous report by King et al., (2023) and summarised our 6 cardio vascular measures into 3 related latent variables. LVF1 captured mainly static blood pressure (BP). LVF2 mainly loaded mainly on pulse pressure (PP). LVF3 loaded on both high and low frequency heart rate

#### Genetic effects on fluid intelligence

The PGS scores were not associated with any of the WM factors (see Sup. Fig. 2). IQ PGS was significantly related to Fluid Intelligence, as expected ($\beta = 0.17$ $t = 5.78$, $p < 0.001$, $\eta^2 = 0.06$). When we controlled for effect of PGS, most WMFs remained related to cognition (see Supp. Table 5). This suggested that the WMFs are capturing brain-cognition associations not fully explained by chronological age or PGS scores for intelligence, suggesting other environmental factors.

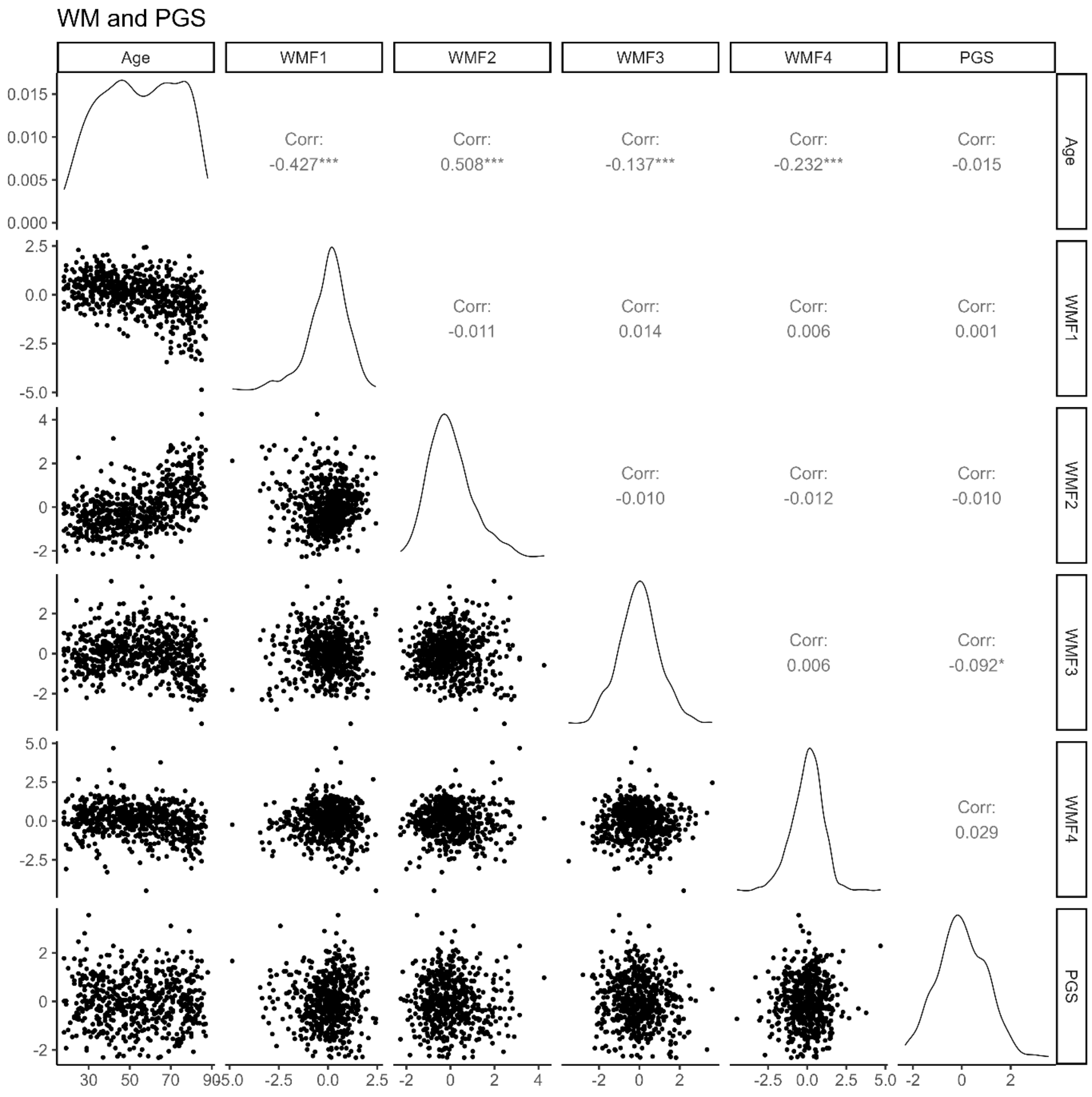


**Supplementary Figure 2** *Correlation matrix between age, each of the 4 WM factors and PGS for cognitive ability.* Matrix shows PGS was not strongly related to any of the WM factors.

#### Cardiovascular and WM factors

For all path analyses we allowed the dependent variables to be correlated.  This meant that covariance between IQ, PS and Memory was accounted for in the model, allowing us to examine unique paths between variables.

**Supplementary Table 1** *Relationship between cardiovascular factors and white matter without accounting for Age.* Table shows estimated direct paths between vascular and white matter factor scores. We found that only pulse pressure factor (LVF2) was significantly related to the white matter factors representing free water content (WMF2).

| lhs | op | rhs | est | se | *z* | *p* | ci.lower | ci.upper |
|---|---|---|---|---|---|---|---|---|
| WMF1 | ~ | LVF1 | 0.00 | 0.04 | 0.11 | .910 | -0.08 | 0.09 |
| WMF1 | ~ | LVF2 | -0.15 | 0.04 | -3.44 | .001*** | -0.24 | -0.07 |
| WMF1 | ~ | LVF3 | 0.22 | 0.04 | 5.51 | < .001*** | 0.14 | 0.30 |
| WMF2 | ~ | LVF1 | -0.03 | 0.04 | -0.77 | .440 | -0.12 | 0.05 |
| WMF2 | ~ | LVF2 | 0.14 | 0.04 | 3.15 | .002** | 0.05 | 0.23 |
| WMF2 | ~ | LVF3 | -0.30 | 0.04 | -7.43 | < .001*** | -0.37 | -0.22 |
| WMF3 | ~ | LVF1 | 0.16 | 0.05 | 3.52 | < .001*** | 0.07 | 0.26 |
| WMF3 | ~ | LVF2 | -0.21 | 0.05 | -4.45 | < .001*** | -0.30 | -0.12 |
| WMF3 | ~ | LVF3 | 0.06 | 0.04 | 1.51 | .131 | -0.02 | 0.15 |
| WMF4 | ~ | LVF1 | 0.06 | 0.05 | 1.21 | .226 | -0.04 | 0.15 |
| WMF4 | ~ | LVF2 | -0.17 | 0.05 | -3.53 | < .001*** | -0.26 | -0.07 |
| WMF4 | ~ | LVF3 | 0.11 | 0.04 | 2.70 | .007** | 0.03 | 0.20 |

#### Cardiovascular and WM factors

**Supplementary Table 2** *Relationship between cardiovascular factors and white matter.* Table shows estimated direct paths between vascular and white matter factor scores. We found that only pulse pressure factor (LVF2) was significantly related to the white matter factors representing free water content (WMF2).

| lhs | op | rhs | est | se | *z* | *p* | ci.lower | ci.upper |
|---|---|---|---|---|---|---|---|---|
| WMF1 | ~ | LVF1 | -0.04 | 0.04 | -0.98 | .325 | -0.12 | 0.04 |
| WMF1 | ~ | LVF2 | -0.03 | 0.05 | -0.61 | .540 | -0.12 | 0.06 |
| WMF1 | ~ | LVF3 | 0.01 | 0.04 | 0.17 | .864 | -0.08 | 0.09 |
| WMF2 | ~ | LVF1 | 0.01 | 0.04 | 0.22 | .830 | -0.07 | 0.09 |
| WMF2 | ~ | LVF2 | -0.10 | 0.04 | -2.31 | .021* | -0.19 | -0.02 |
| WMF2 | ~ | LVF3 | -0.02 | 0.04 | -0.55 | .585 | -0.11 | 0.06 |
| WMF3 | ~ | LVF1 | 0.09 | 0.05 | 1.85 | .064 | -0.01 | 0.18 |
| WMF3 | ~ | LVF2 | -0.10 | 0.05 | -1.96 | .050* | -0.20 | -0.00 |
| WMF3 | ~ | LVF3 | -0.02 | 0.05 | -0.44 | .662 | -0.12 | 0.07 |
| WMF4 | ~ | LVF1 | 0.02 | 0.05 | 0.51 | .609 | -0.07 | 0.11 |
| WMF4 | ~ | LVF2 | -0.01 | 0.05 | -0.25 | .804 | -0.11 | 0.09 |
| WMF4 | ~ | LVF3 | 0.00 | 0.05 | 0.03 | .977 | -0.09 | 0.10 |
| WMF1 | ~ | Age | -10.39 | 1.27 | -8.18 | < .001*** | -12.88 | -7.90 |
| WMF1 | ~ | QuadAge | -3.62 | 0.99 | -3.65 | < .001*** | -5.56 | -1.68 |
| WMF1 | ~ | Sex | -0.19 | 0.03 | -5.35 | < .001*** | -0.25 | -0.12 |
| WMF2 | ~ | Age | 14.25 | 1.23 | 11.61 | < .001*** | 11.84 | 16.65 |
| WMF2 | ~ | QuadAge | 5.68 | 0.96 | 5.94 | < .001*** | 3.81 | 7.55 |
| WMF2 | ~ | Sex | -0.14 | 0.03 | -4.32 | < .001*** | -0.21 | -0.08 |
| WMF3 | ~ | Age | -3.90 | 1.40 | -2.79 | .005** | -6.64 | -1.16 |
| WMF3 | ~ | QuadAge | -7.21 | 1.09 | -6.61 | < .001*** | -9.35 | -5.07 |
| WMF3 | ~ | Sex | -0.09 | 0.04 | -2.33 | .020* | -0.16 | -0.01 |
| WMF4 | ~ | Age | -6.15 | 1.41 | -4.38 | < .001*** | -8.90 | -3.40 |
| WMF4 | ~ | QuadAge | -4.85 | 1.10 | -4.43 | < .001*** | -7.00 | -2.71 |
| WMF4 | ~ | Sex | 0.20 | 0.04 | 5.32 | < .001*** | 0.13 | 0.28 |

#### WM and Cognitive factors not controlling for age

**Supplementary Table 3** *Relationship between white matter factors and cognition without accounting for Age and Sex.* Table shows estimated direct paths between white matter factor scores and cognitive scores. We found that all paths between our white matter factors and cognitive measures were significant once we did not correct for age and sex effects.

| lhs | op | rhs | est | se | *z* | *p* | ci.lower | ci.upper |
|---|---|---|---|---|---|---|---|---|
| IQ | ~ | WMF1 | 0.38 | 0.03 | 11.46 | < .001*** | 0.32 | 0.45 |
| IQ | ~ | WMF2 | -0.35 | 0.03 | -10.65 | < .001*** | -0.41 | -0.28 |
| IQ | ~ | WMF3 | 0.10 | 0.03 | 3.02 | .003** | 0.03 | 0.16 |
| IQ | ~ | WMF4 | 0.24 | 0.03 | 7.30 | < .001*** | 0.17 | 0.30 |
| PS | ~ | WMF1 | 0.41 | 0.03 | 12.41 | < .001*** | 0.34 | 0.47 |
| PS | ~ | WMF2 | -0.35 | 0.03 | -10.57 | < .001*** | -0.41 | -0.28 |
| PS | ~ | WMF3 | 0.07 | 0.03 | 2.17 | .030* | 0.01 | 0.14 |
| PS | ~ | WMF4 | 0.17 | 0.03 | 5.15 | < .001*** | 0.11 | 0.24 |
| Mem | ~ | WMF1 | 0.19 | 0.04 | 5.03 | < .001*** | 0.11 | 0.26 |
| Mem | ~ | WMF2 | -0.20 | 0.04 | -5.39 | < .001*** | -0.27 | -0.13 |
| Mem | ~ | WMF3 | 0.08 | 0.04 | 2.22 | .026* | 0.01 | 0.15 |
| Mem | ~ | WMF4 | 0.15 | 0.04 | 4.05 | < .001*** | 0.08 | 0.22 |

#### WM and Cognitive factors controlling for age

**Supplementary Table 4** *Relationship between white matter factors and cognition.* Table shows estimated direct paths between white matter factor scores and cognitive scores. Fluid intelligence (IQ) was predicted by the first and fourth white matter factors capturing measures of white matter microstructure. Processing speed (PS) was predicted mainly by Factor 1, and showed a less robust association with Factor 2, representing free water content. Episodic Memory (Mem) was not significantly related to any of the white matter factors after accounting for age and sex effects.

| lhs | op | rhs | est | se | *z* | *p* | ci.lower | ci.upper |
|---|---|---|---|---|---|---|---|---|
| IQ | ~ | WMF1 | 0.09 | 0.04 | 2.49 | .013* | 0.02 | 0.16 |
| IQ | ~ | WMF2 | -0.03 | 0.04 | -0.88 | .379 | -0.11 | 0.04 |
| IQ | ~ | WMF3 | -0.02 | 0.03 | -0.69 | .488 | -0.08 | 0.04 |
| IQ | ~ | WMF4 | 0.10 | 0.03 | 3.02 | .003** | 0.03 | 0.16 |
| PS | ~ | WMF1 | 0.17 | 0.04 | 4.74 | < .001*** | 0.10 | 0.25 |
| PS | ~ | WMF2 | -0.08 | 0.04 | -1.99 | .047* | -0.16 | -0.00 |
| PS | ~ | WMF3 | -0.01 | 0.03 | -0.26 | .799 | -0.07 | 0.06 |
| PS | ~ | WMF4 | 0.06 | 0.03 | 1.79 | .074 | -0.01 | 0.12 |
| Mem | ~ | WMF1 | 0.04 | 0.04 | 1.00 | .318 | -0.04 | 0.13 |
| Mem | ~ | WMF2 | 0.02 | 0.05 | 0.44 | .663 | -0.07 | 0.11 |
| Mem | ~ | WMF3 | 0.02 | 0.04 | 0.64 | .522 | -0.05 | 0.10 |
| Mem | ~ | WMF4 | 0.03 | 0.04 | 0.63 | .528 | -0.05 | 0.10 |
| IQ | ~ | Age | -16.09 | 1.20 | -13.40 | < .001*** | -18.44 | -13.73 |
| IQ | ~ | QuadAge | -3.74 | 0.94 | -3.97 | < .001*** | -5.59 | -1.89 |
| IQ | ~ | Sex | -0.09 | 0.03 | -2.88 | .004** | -0.15 | -0.03 |
| PS | ~ | Age | -14.40 | 1.23 | -11.67 | < .001*** | -16.82 | -11.98 |
| PS | ~ | QuadAge | -0.63 | 0.95 | -0.67 | .504 | -2.49 | 1.22 |
| PS | ~ | Sex | -0.05 | 0.03 | -1.50 | .134 | -0.11 | 0.01 |
| Mem | ~ | Age | -9.99 | 1.47 | -6.80 | < .001*** | -12.87 | -7.11 |
| Mem | ~ | QuadAge | -2.03 | 1.15 | -1.77 | .077 | -4.27 | 0.22 |
| Mem | ~ | Sex | 0.14 | 0.04 | 3.80 | < .001*** | 0.07 | 0.21 |

#### WM and Cognitive factors controlling for age and polygenic effects.

**Supplementary Table 5** *Relationship between white matter factors and cognition controlling for polygenic effects.* Table shows estimated direct paths between WMFs and cognitive scores after accounting for age and polygenic effects. Fluid intelligence (IQ) was predicted by the first and fourth WMFs capturing measures of white matter microstructure. Processing speed (PS) was predicted mainly by Factor 1, but was not associated with Factor 2. Episodic Memory (Mem) was not significantly related to any of the WMFs after accounting for age and sex effects.

| lhs | op | rhs | est | se | *z* | *p* | ci.lower | ci.upper |
|---|---|---|---|---|---|---|---|---|
| IQ | ~ | WMF1 | 0.09 | 0.04 | 2.53 | .011* | 0.02 | 0.17 |
| IQ | ~ | WMF2 | -0.02 | 0.04 | -0.53 | .597 | -0.10 | 0.06 |
| IQ | ~ | WMF3 | 0.00 | 0.03 | 0.00 | .998 | -0.07 | 0.07 |
| IQ | ~ | WMF4 | 0.09 | 0.03 | 2.88 | .004** | 0.03 | 0.16 |
| PS | ~ | WMF1 | 0.15 | 0.04 | 3.95 | < .001*** | 0.08 | 0.23 |
| PS | ~ | WMF2 | -0.04 | 0.04 | -0.95 | .340 | -0.12 | 0.04 |
| PS | ~ | WMF3 | 0.02 | 0.03 | 0.61 | .540 | -0.05 | 0.09 |
| PS | ~ | WMF4 | 0.05 | 0.03 | 1.48 | .139 | -0.02 | 0.12 |
| Mem | ~ | WMF1 | 0.05 | 0.05 | 1.07 | .286 | -0.04 | 0.15 |
| Mem | ~ | WMF2 | 0.01 | 0.05 | 0.17 | .864 | -0.09 | 0.11 |
| Mem | ~ | WMF3 | 0.06 | 0.04 | 1.38 | .167 | -0.03 | 0.14 |
| Mem | ~ | WMF4 | 0.02 | 0.04 | 0.52 | .606 | -0.06 | 0.11 |
| IQ | ~ | Age | -15.90 | 1.27 | -12.51 | < .001*** | -18.39 | -13.41 |
| IQ | ~ | QuadAge | -3.52 | 0.98 | -3.60 | < .001*** | -5.44 | -1.61 |
| IQ | ~ | Sex | -0.08 | 0.03 | -2.56 | .010* | -0.14 | -0.02 |
| PS | ~ | Age | -15.56 | 1.31 | -11.88 | < .001*** | -18.13 | -13.00 |
| PS | ~ | QuadAge | -0.44 | 1.01 | -0.44 | .660 | -2.42 | 1.53 |
| PS | ~ | Sex | -0.03 | 0.03 | -0.86 | .390 | -0.09 | 0.04 |
| Mem | ~ | Age | -8.66 | 1.66 | -5.22 | < .001*** | -11.90 | -5.41 |
| Mem | ~ | QuadAge | -0.66 | 1.28 | -0.52 | .604 | -3.16 | 1.84 |
| Mem | ~ | Sex | 0.15 | 0.04 | 3.70 | < .001*** | 0.07 | 0.23 |
| IQ | ~ | PGS | 0.15 | 0.03 | 5.16 | < .001*** | 0.10 | 0.21 |
| PS | ~ | PGS | 0.11 | 0.03 | 3.47 | .001*** | 0.05 | 0.17 |
| Mem | ~ | PGS | 0.17 | 0.04 | 4.42 | < .001*** | 0.10 | 0.25 |

#### Individual white measures and Cognition.

We examined how individual WM measures predicted cognition. To make sure the comparison is fair, we focused on the 570 participants that had data for all 11 WM measures. We predicted each individual cognitive measure from each individual WM measure while controlling for age and sex (DV ~ WM_measure + agepoly_1 + agepoly_2 + Sex). The results could guide future neurocognitive studies of ageing when choosing a specific measure of WM.

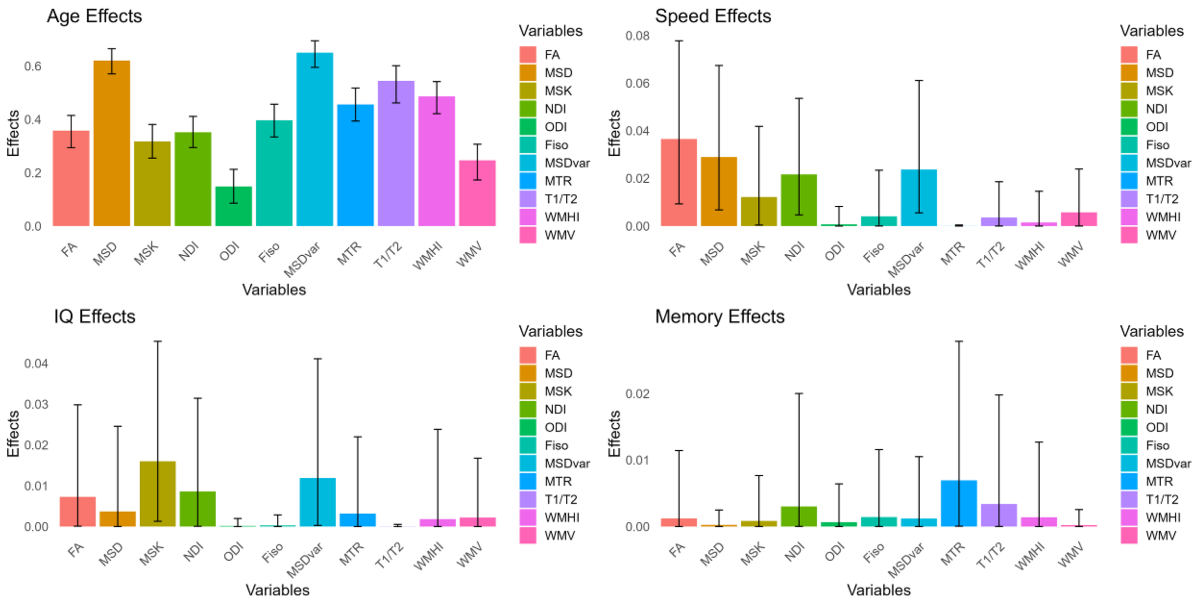


**Supplementary Figure 3** *Effect sizes for each individual WM measure and age and cognition.* We show partial eta squared for each measure predicting each of the cognitive measures and cognition independently. We used 1000 bootstraps to show confidence interval over the effect size. As indicated by the factor analysis, kurtosis is strongest predictor of fluid intelligence, whereas FA, MSD and NDI are stronger predictors for processing speed.

Similarly we show how well each of the latent cardiovascular factors predicted each individual WM measure (WM ~  LVF1 + LVF2 + LVF3 + agepoly_1 + agepoly_2 + Sex).

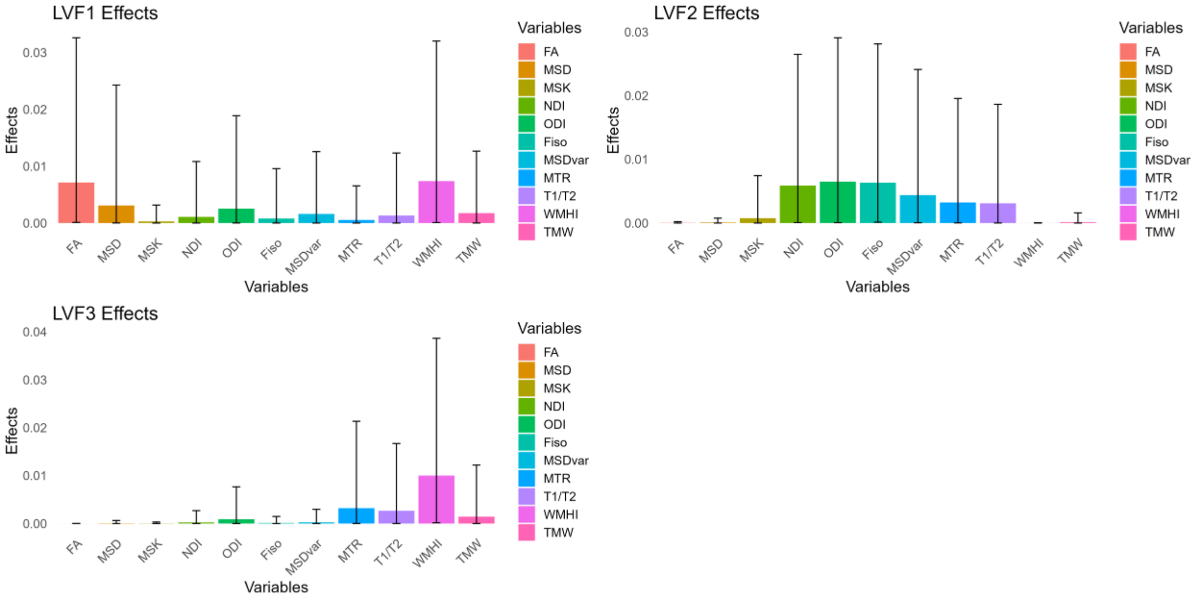


**Supplementary Figure 4** Figure shows how strongly each latent cardiovascular factor is related to each individual white matter measure. We show partial eta square and bootstrapped confidence intervals.

**Supplementary Table 6. Effect size of each WM measure.** Table shows partial eta square for independent regressions for each individual WM measure predicting Age, Fluid intelligence (IQ), Processing Speed (PS) and Episodic Memory (Mem). See also Supplementary figure 3.

| Measure | Age | IQ | PS | Mem |
|---|---|---|---|---|
| FA | 0.36 | 0.01 | 0.04 | 0.00 |
| MSD | 0.62 | 0.00 | 0.03 | 0.00 |
| MSK | 0.32 | 0.02 | 0.01 | 0.00 |
| NDI | 0.35 | 0.01 | 0.02 | 0.00 |
| ODI | 0.15 | 0.00 | 0.00 | 0.00 |
| Fiso | 0.40 | 0.00 | 0.00 | 0.00 |
| MSDvar | 0.65 | 0.01 | 0.02 | 0.00 |
| MTR | 0.46 | 0.00 | 0.00 | 0.01 |
| T1/T2 | 0.54 | 0.00 | 0.00 | 0.00 |
| WMHI | 0.49 | 0.00 | 0.00 | 0.00 |
| WMV | 0.25 | 0.00 | 0.01 | 0.00 |

**Supplementary Table 7 Effect size of each cardiovascular measure.** Table shows partial eta square for regression predicting each individual WM measure by our 3 latent cardiovascular factors and age and sex confounds. See also Supplementary figure 4.

| Measure | LVF1 | LVF2 | LVF3 |
|---|---|---|---|
| FA | 0.01 | 0.00 | 0.00 |
| MSD | 0.00 | 0.00 | 0.00 |
| MSK | 0.00 | 0.00 | 0.00 |
| NDI | 0.00 | 0.01 | 0.00 |
| ODI | 0.00 | 0.01 | 0.00 |
| Fiso | 0.00 | 0.01 | 0.00 |
| MSDvar | 0.00 | 0.00 | 0.00 |
| MTR | 0.00 | 0.00 | 0.00 |
| T1/T2 | 0.00 | 0.00 | 0.00 |
| WMHI | 0.01 | 0.00 | 0.01 |
| WMV | 0.00 | 0.00 | 0.00 |

#### Factor analysis for only diffusion metrics

**Supplementary Table 8** *PCA for DWI metrics*. This table shows number of components identified for the same PCA analyses where we only used the 6 DWI measures. We show the variance explained (VE) by the last component (Last Comp) and the cumulative variance explained by the number of components retained.

| Measures | N Measures | N Comp *ekf* | VE Last Comp | Cum VE |
|---|---|---|---|---|
| DWI | 6 | 3 | 11.63 | 92.44 |
